# Topic Modeling analysis of the Allen Human Brain Atlas

**DOI:** 10.1101/2024.10.11.617855

**Authors:** Letizia Pizzini, Filippo Valle, Matteo Osella, Michele Caselle

**Affiliations:** Department of Physics and INFN, University of Turin, via P.Giuria 1, 10125 Turin, Italy

**Keywords:** Human Brain, Stochastic Block Modeling, Network Theory, Topic Modeling, Spatial Transcriptomics, Gene Expression

## Abstract

The human brain is a complex interconnected structure controlling all elementary and high-level cognitive tasks. It is composed of many regions that exhibit specific distributions of cell types and distinct patterns of functional connections. This complexity is rooted in differential transcription. The constituent cell types of different brain regions express distinctive combinations of genes as they develop and mature, ultimately shaping their functional state in adulthood. How precisely the genetic information of anatomical structures is connected to their underlying biological functions remains an open question in modern neuroscience.

A major challenge is the identification of “universal patterns”, which do not depend on the particular individual, but are instead basic structural properties shared by all brains. Despite the vast amount of gene expression data available at both the bulk and single-cell levels, this task remains challenging, mainly due to the lack of suitable data mining tools.

In this paper, we propose an approach to address this issue based on a hierarchical version of Stochastic Block Modeling. Thanks to its specific choice of priors, the method is particularly effective in identifying these universal features. We use as a laboratory to test our algorithm a dataset obtained from six independent human brains from the Allen Human Brain Atlas. We show that the proposed method is indeed able to identify universal patterns much better than more traditional algorithms such as Latent Dirichlet Allocation or Weighted Correlation Network Analysis. The probabilistic association between genes and samples that we find well represents the known anatomical and functional brain organization. Moreover, leveraging the peculiar “fuzzy” structure of the gene sets obtained with our method, we identify examples of transcriptional and post-transcriptional pathways associated with specific brain regions, highlighting the potential of our approach.

## 1 Introduction

Over the past two decades, the analysis of transcriptomic data has emerged as a powerful approach for understanding the molecular foundation of large-scale neural characteristics observed throughout the entire brain. While in the past technical limitations constrained analyses of gene expression in the brain to small sets of areas studied in isolation, in the last few years, the impressive advances in gene expression technologies allowed genome-wide measurements of gene expression and chromatin accessibility both in bulk tissue and at the single-cell level^1,2^. Datasets with millions of single-cell data for several different cell populations are now available^3,4^. This vast amount of information requires new, innovative data-mining tools. A significant challenge presented by this data is the presence of inter-individual differences, which often prevent the identification of universal behaviors common to all individuals. This contrasts with the typical analysis of other tissues, and it is probably a consequence of the huge complexity of the brain. However, it is exactly this interplay of complex but common interactions that we would like to extract from genomic data of the brain. It is thus important to devise data mining tools able to overcome the inter-individual differences. Most of the existing tools^5^, as we shall see below, are based on statistical priors or, more generally, on pre-processing strategies that go exactly in the opposite direction. In fact, they try to amplify the differences between groups of samples in order to identify “marker genes” of a given tissue or condition. The main goal of the present paper is to discuss a computational framework which avoids this bias, and to show that it indeed allows for the extraction of universal features. To test this issue in a controlled way, we used as a laboratory a well-studied dataset from the Allen Human Brain Atlas consortium^6^. This is a well established and highly curated dataset that has been studied in several papers over the last few years^7–9^.

The Allen Human Brain Atlas (AHBA) https://human.brain-map.org/ provides detailed and comprehensive coverage of almost the entire brain, quantifying the expression level of more than 20 000 genes in 3 702 spatially distinct tissue samples taken from six neurotypical adult brains. This dataset represented a relevant achievement in the field and the starting point for a wealth of important follow-up studies^10–12^. At the same time, it represents a perfect example of the challenge discussed above. The data is characterized by significant inter-individual differences in transcriptional profiles^8,13,14^ related to different ethnic backgrounds, genders, medical histories, causes of death, ages, and varying post-mortem intervals of the six individuals (see details in Table 1 in the Supplementary Material). As a consequence, it often happens that samples from different regions of the same brain show more similar gene expression patterns than samples taken from the same location in different brains. This makes the AHBA dataset a perfect laboratory to test our tools, with the aim of applying them in the future to the richer and more complex datasets now available^1–4^.

**Table 1.** Gene Set Enrichment Analysis for different topics. . The number of genes in each topic is shown in square brackets. The analysis was only conducted for those topics composed of a sufficiently high number of genes. For each topic, only the sets that have a higher score based on the FDR q-Value are reported. Many of our topics are enriched for annotated gene sets related to the brain structures associated with them through the analysis in Figure 3. Others are linked to terms related to brain functions that are commonly associated with the identified regions.

| hSBM Topic [# Genes] | Gene Set Name | FDR q-Value |
| --- | --- | --- |
| Topic 2 [112] | GOCC.SYNAPSE | 5.44e<sup>-27</sup> |
|  | GOCC.SYNAPTIC_MEMBRANE | 6.44e<sup>-22</sup> |
|  | GOCC.NEURON_PROJECTION | 9.74e<sup>-22</sup> |
|  | GOBP.MEMORY | 1.36e<sup>-8</sup> |
| Topic 3 [110] | LEIN_OLIGODENDROCYTE_MARKERS | 1.65e<sup>-16</sup> |
|  | GOBP.ENSHEATHMENT_OF_NEURONS | 3.61e<sup>-11</sup> |
|  | GOBERT_OLIGODENDROCYTE_DIFFERENTIATION_ION_DN | 3.66e<sup>-10</sup> |
|  | GOBP.CENTRAL_NERVOUS_SYSTEM_DEVELOPMENT | 9.97e<sup>-10</sup> |
| Topic 5 [89] | GOCC.SYNAPSE | 1.39e<sup>-19</sup> |
|  | GOCC.SYNAPTIC_MEMBRANE | 3.04e<sup>-18</sup> |
|  | GOCC.POSTSYNAPSE | 8.24e<sup>-17</sup> |
|  | REACTOME.NEURONAL_SYSTEM | 5.9e<sup>-14</sup> |
| Topic 9 [77] | GOBP.NERVOUS_SYSTEM_PROCESS | 1.23e<sup>-9</sup> |
|  | GOBP.BEHAVIOR | 3.5e<sup>-6</sup> |
|  | GOBP.RESPONSE_TO_ENDOGENOUS_STIMULUS | 3.5e<sup>-6</sup> |
|  | GOBP.LOCOMOTORY_BEHAVIOR | 5.1e<sup>-6</sup> |
| Topic 11 [66] | LEIN.CEREBELLUM_MARKERS | 1.5e<sup>-8</sup> |
| Topic 13 [49] | GOBP.CELL_CELL_SIGNALING | 1.66e<sup>-8</sup> |
|  | GOBP.SYNAPTIC_SIGNALING | 1.66e<sup>-8</sup> |
| Topic 19 [30] | BLALOCK_ALZHEIMERS_DISEASE_DN | 1.89e<sup>-8</sup> |
|  | STARK_PREFRONTAL_CORTEX_22Q11_DELETION_UP | 7.67e<sup>-7</sup> |
|  | KIM_ALL_DISORDERS_CALB1_CORR_UP | 7.67e<sup>-7</sup> |
|  | LU_AGING_BRAIN_DN | 4.99e<sup>-6</sup> |
| Topic 20 [24] | ASTON_MAJOR_DEPRESSIVE_DISORDERS_DN | 7.54e<sup>-6</sup> |
|  | WP_CELL_LINEAGE_MAP_FOR_NEURONAL_DIFFERENTIATION | 3.83e<sup>-5</sup> |

The usual approach to address the issue of inter-individual differences is to impose an a-priori selection of genes showing the same behavior across subjects^9^, which, however, may lead to a significant loss of information.

The main goal of our paper is to propose a different strategy to address this problem. Specifically we will use a new class of clustering algorithms that represent an alternative approach to topic modelling based on a hierarchical version of the Stochastic Block Modelling (hSBM)^15^, which we previously introduced in the the context of cancer genomics^16^.

Topic modeling algorithms were proposed a few years ago^17^ in the context of Natural Language Processing (NLP) to infer the latent topical structure of a collection of documents. Then, in the past few years, their application has been also extended to transcriptomic data. In particular, they have been successfully used to perform unsupervised clustering in the gene expression space^16,18–20^. The idea of exploiting computational methods born to be used in the field of text mining is based on the observation that text mining systems face similar challenges in terms of complexity and heterogeneity as the problem of gene expression in tissue samples^21^. Topic modeling algorithms try to discover the topics present in a corpus of texts by analyzing word frequencies. In our context, tissue samples correspond to documents, genes represent words, and the frequency of word usage in a document is analogous to the expression level of a gene in a sample. In both cases, the goal is to cluster together samples (or documents) that deal with the same gene sets (topics) and, specularly, obtain the genetic signatures of specific groups of samples involved in the same structures or common functional patterns.

What substantially differentiates topic modeling tools from other classical clustering approaches is their probabilistic nature, which provides much more information than standard clustering algorithms. Topic modeling algorithms allow not only to cluster samples and/or genes but also return the probability of each gene to characterize a topic and the probability of a sample to be described by a given topic. In our case, this will allow us to detect non trivial associations between brain regions (or other types of sample clusters) and groups of genes (topics).

The most popular topic modeling algorithm is the so-called Latent Dirichlet Allocation (LDA) algorithm^17^, which assumes a Dirichlet prior for the topics distribution. This assumption finds no basis or justification either in textual problems nor in biological systems, but makes the algorithm very efficient even when applied to large databases.

In recent years, significant progress has been made in the field, which has set the groundwork for addressing and surpassing the limitations of LDA, such as the lack of biological rationale behind the prior and the presence of many free parameters. A valid alternative comes from the adaptation of community detection techniques^22^, widely studied in network theory, for the purpose of topic modeling^15^. This type of algorithm, based on hierarchical Stochastic Block Model (hSBM), does not require assumptions on the probability distribution of the latent variables, and no parameter must be set a priori, in order to successfully handle the analysis of heterogeneous and complex systems such as the classification of tumor subtypes based on gene expression data^16^, multiomic data^19^, and even single-cell gene expression data^20^.

hSBM reformulates topic modeling as a community detection problem in a bipartite network where words (genes) belong to the first layer and documents (tissue samples) to the second one. The algorithm is based on a flat prior, which avoids the problems of LDA, and is structured as a hierarchy of nested SBMs, leading to a hierarchical organization of samples (and genes). The algorithm autonomously returns the optimal number of topics and clusters for each hierarchical level with no need for external input or hyperparameter tuning.

As we shall see in detail below, hSBM shows remarkable robustness with respect to gene selection choices and to the pre-processing protocol. Moreover, the peculiar organization of the output in terms of the probability distribution of genes over topics and topics over samples allows for a sort of “fuzzy” clustering of genes, which is better suited than standard clustering algorithms to describe the variety of different cell populations in the brain. To support this observation, we shall compare hierarchical Stochastic Block Model (hSBM)^15^ results with those of other popular algorithms like Latent Dirichlet Allocation (LDA)^17^ and Weighted Gene Correlation Network Analysis (WGCNA)^23^. As we shall see, hSBM outperforms the other algorithms both in identifying meaningful biological annotations and in finding “universal” features, conserved across different subjects, which is the main goal of our work.

## 2 Results

### 2.1 A topic modeling analysis of the AHBA dataset based on hSBM

The hierarchical Stochastic Block Model (hSBM)^15,24^ belongs to the class of probabilistic inference approaches to community detection in bipartite networks. The model makes minimal assumptions on the data, structure compared to competing algorithms, and it is able to directly infer the hierarchical network structure. In other words, it automatically detects the number of topics and hierarchically clusters both sides of the network (in our case, both genes and samples), thus overcoming some common limitations of standard clustering approaches^16^.

These properties make this algorithm a perfect tool to address gene expression data which are highly heterogeneous and organized in hierarchical structures with unknown number of layers and clusters. This is particularly true for the AHBA dataset, which, as we shall see below, has a well-defined hierarchical organization of the samples.

The starting point of our analysis is to represent the AHBA data as a bipartite network. In principle, we would expect to have a network relating genes and samples (this is typically the case when dealing with RNA-sequencing data). However, the actual entries of the AHBA are microarray *probes* and, generally, a gene is associated with more than one probe. For reasons which we will clarify in detail later, we decided to use the probes and not the genes as nodes of the gene layer of the bipartite network. Therefore, the network is composed by links connecting probes {*p_i_*} with samples {*s_j_*}. Each link has a weight *w_i j_* that encodes the expression level of the probe *p_i_* in the sample *s _j_*. Each probe can then be associated to the corresponding gene when the results are analyzed and interpreted.

The hSBM output provides a hierarchical and probabilistic structure of the data into “blocks”. Samples and probes have an associated probability of belonging to a particular block (see the description of hSBM in the Methods section). We define the blocks of samples as *clusters* and the blocks of probes as *topics* (Fig. 1). By construction, probes are included in topics (and samples in clusters) with a hard membership: each probe is associated only to a given topic and each sample only to a given cluster. However, even if we have a hard membership at the level of probes, we have a fuzzy membership in the gene space. This happens when different probes of the same gene fall into different topics. In this case, we associate the corresponding gene to all these topics. This may happen due to alternative splicing or when the gene is effectively in an intermediate position between the two topics and stochastic fluctuations of the expression levels of the probes are sufficient to create this mixed membership. This event is not frequent but it is still relevant for the analysis. Few relevant examples of this mixed memebership will be discussed in the following sections.

**Figure 1.**
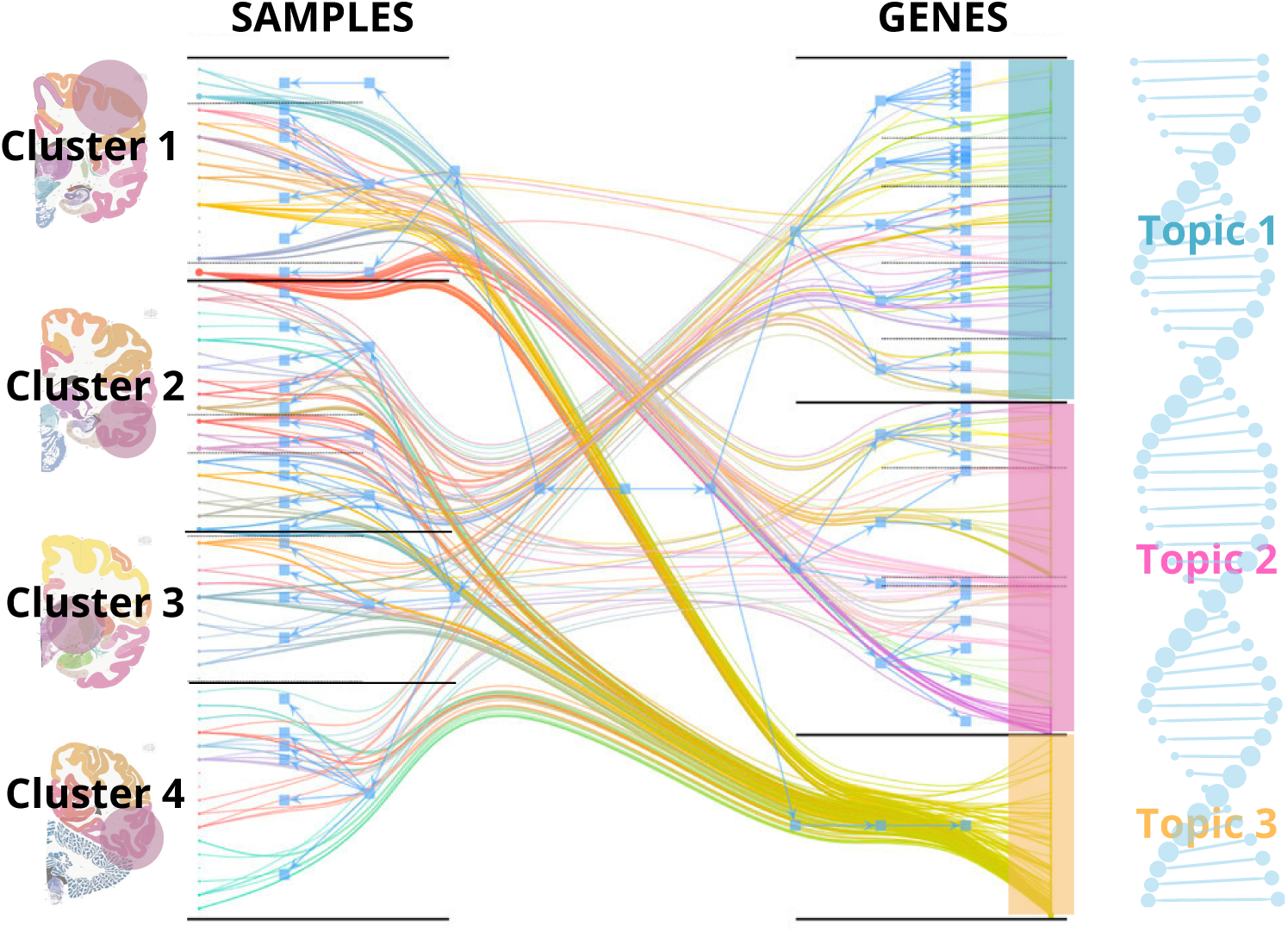
Gene expression data as a bipartite network on which hierarchical stochastic block model (hSBM) organizes genes into topics and samples into clusters. The lines connecting genes and samples encode the weights of the bipartite network (i.e., the gene expression values in the different samples).

As a result of the hSBM clustering, the whole information content of the database is eventually encoded in the two probability distributions *P*(topic*|*sample) and *P*(probe*|*topic) (see the methods section for a precise definition). At this point each probe is associated to the corresponding gene and we can obtain from *P*(probe*|*topic) the corresponding *P*(gene*|*topic). Let us discuss in detail these two probability distributions.

- **P**(**topic***|***sample**) represents the probability that a sample’s gene expression pattern is controlled by the cooperative interaction of a specific set of genes (i.e., a topic). These probabilities follow a ‘many-to-many’ membership structure. In other words, a single sample can be regulated by multiple gene sets (topics), and a particular gene set can affect various samples, potentially across different clusters, which may correspond to distinct brain regions.
- **P**(**gene***|***topic**) depicts the probability distribution of genes within a topic. Not all genes in a topic hold equal importance. Their assignment is weighted according to the *P*(gene*|*topic) distribution. This weighting helps identifing the key genes — those with higher *P*(gene|topic) values — that play the most significant role and are likely driving the gene set’s influence on the samples.

These probability distributions will be the main tool that we will use in the following to extract information from the dataset.

#### 2.1.1 Comparing different clustering algorithms

A major goal of our analysis is to demonstrate that extracting meaningful biological information from a complex dataset like AHBA requires considering the combined effect of the entire set of genes, rather than relying solely on the behavior of isolated markers of a given brain structure or cellular population. This is the same strategy we adopted when addressing cancer expression datasets with the goal of identifying the cancer subtype organization^16,19^. To quantify the efficacy of this strategy, we will compare the performance of hSBM with those of a set of other state-of-the-art clustering tools: Weighted Gene Correlation Network Analysis (WGCNA)^23^, Latent Dirichlet Allocation (LDA)^17^ and hierarchical clustering (hierarchical)^25^.

### 2.2 Clustering of the AHBA dataset

As we mentioned above the AHBA dataset can be represented as a bipartite network with samples on one side and genes on the other side. Both sides present nontrivial features which must be addressed before using any clustering algorithm. Let us see these issue in detail.

#### 2.2.1 The sample side: combining together different individuals

The AHBA offers gene expression data for multiple spatially localized tissue samples, with a total of 232 distinct brain structures being sampled at least once in at least one brain. The Methods section reports a detailed description. These structures are organized in a hierarchical way: each sample is associated with a unique anatomical structure ID, which can be linked to corresponding higher-order structures using the Allen Institute anatomical ontology^8^. This enables the identification of brain structures at different scales of resolution, a feature that will play a major role in the following, since also the results of the topic modeling analysis show a similar hierarchical organization. In particular, following^9^ we concentrate on two levels of organization: a “coarse” one in which the brain is divided into 8 structures that we shall denote in the following as “regions”, and a second level composed by 24 structures that we denote as “subregions”. Table S1 in the Supplementary Materials report a more detailed description of these structures.

Following^9^ we combine together all the samples in a unique large dataset composed by 3 702 samples annotated at the different hierarchical levels mentioned above. The samples correspond to six different individuals and one of our goals is to find *universal* features that are in common among the six individuals. As we mentioned above, despite the remarkable curation of the dataset**^?^**^,9^, the gene expression profiles are still characterized by relevant inter-individual differences^8^. Threfore, a major challenge for our algorithm (and in fact for any clustering algorithm) will be to extract universal features despite the persistent individual bias.

#### 2.2.2 The gene side: filtering and pre-processing

On the gene side, the AHBA consists of more than 58 000 probes associated with more than 20 000 unique protein-coding genes, of which only a portion is presumed to exhibit consistent regional variation in expression across brain structures. Thus, gene filtering is essential to reduce the computational cost of the employed algorithms. The standard approach in the analysis of gene expression data is to select highly variable genes (highly variable probes in our case) which, being differentially expressed among samples, may improve the performance of clustering algorithms. However, given the nature of the dataset in analysis, this is a rather non-trivial procedure and there are two alternative ways to perform this selection.

The first option is to select probes with the highest variability within the merged dataset of the six brains. The second option is to independently select highly variable probes for each brain and merge the datasets by keeping only the probes common to all selections (see the Methods section for a detailed discussion). This second choice avoids retaining genes that predominantly exhibit expression variation between individual brains, This choice would amplify inter-individual differences within the dataset, but also decreases the probability of selecting biologically significant genes that define cellular distinctions, and thus the structural organization of the brain.

In the following, we compare both strategies. In both cases, we select the 4 000 most variable probes. In the first case, these highly variable probes are associated with 1 210 different genes, while in the second case, they are associated with 1 743 genes. The overlap between the two selections turns out to be only 2 750 probes corresponding to 1 001 genes. The difference between the two sets is thus not negligible and will allow us to test the robustness of hSBM with respect to the gene selection choice.

For sake of brevity we report in the main text only the results obtained with the first choice (highly variable genes derived after merging together the six brains). In the Supplementary Material (Fig. S4), we include a comparison with the results obtained with the second choice, which serves as evidence of the algorithm robustness to the gene selection procedure.

### 2.3 Main features of the hSBM clustering of the AHBA

hSBM identifies a hierarchical structure with four levels of resolution, of which the first and the last are uninformative so we shall neglect them in the following analysis. The first layer divides samples and genes in 2 clusters and 2 topics respectively while in the latter the algorithm finds 158 clusters and 331 topics. The most interesting results are collected in the second and third layers.

In the second layer, the samples are grouped into 9 clusters, and the genes are arranged into 32 topics. From an anatomical point of view, this resolution corresponds to the “region” level of the AHBA, and remarkably, the number of clusters (9) is very similar to the number (8) of regions into which the AHBA samples are divided. The third layer is composed of 41 clusters and 164 topics, and corresponds anatomically to the “subregion” level (the files with the results are made available in the Supplementary section). It is worthwhile to stress that the algorithm identified these two levels of resolution *with no external input*. This is one of the most interesting features of hSBM with respect to more conventional clustering algorithms, which usually require the number of clusters to be given as input. In the following, we will mainly concentrate on the second layer, which is simpler to handle thanks to the small number of regions and topics, and thus allows a more in depth discussion.

We labelled the 32 topics according to their size. The size distribution is reported in Figure S2 of the Supplementary Material. The tail (in red) of the distribution contains topics with 10 or fewer genes, which we excluded, as having fewer than 10 genes makes it difficult to perform subsequent analyses, such as gene set enrichment analysis. We end up with a partition of the genes into 27 topics, which we shall analyze in detail in the following.

As mentioned above, we have a hard membership of topics at the level of probes, but a fuzzy membership in the genes space. To quantify this phenomenon, we report in Figure S3 of the Supplementary Material a heat map in which each off-diagonal entry reports the number of genes in common between the two topics that correspond to the column and row of the entry. This fuzzy membership will play a major role in the following.

### 2.4 The hierarchical organization of hSBM clusters mimics the anatomical organization of the brain

The most significant results obtained from the application of the algorithms to the AHBA dataset are reported in the Figure 2, where we compare the hSBM clustering of samples with the brain partitions of the AHBA.

**Figure 2.**
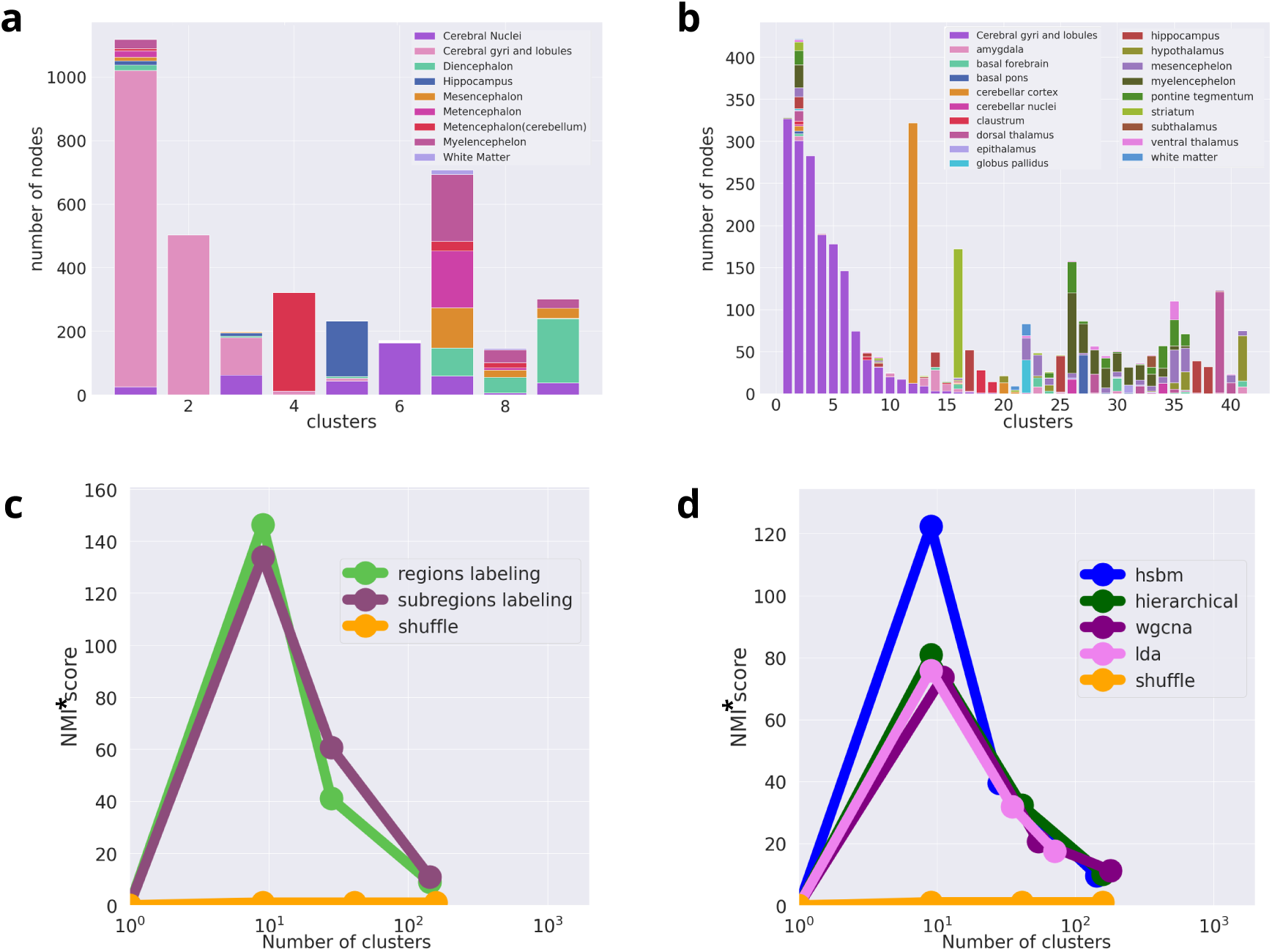
Topic modeling results for the AHBA. Panel (**a**) and (**b**) report the organization of samples in clusters obtained via hSBM. Each column is a cluster while colors represent the different anatomical regions reported in the legend. The height of each column is proportional to the number of samples in the cluster. In (**a**), we report the results for the second layer of clustering (9 clusters) compared with a coarse-grained organization of anatomical regions. (**b**) represents the sample organization at the third layer (41 clusters) compared with a finer ontology of subregions. In (**c**), we show the NMI* scores of the comparison between the hSBM clusters and the anatomical labeling of regions (green dots) and subregions (violet dots). (**d**) The comparison of NMI* scores across the four hierarchy levels is reported for different clustering algorithms, using the anatomical labeling of the brain regions.

Figure 2**a**,**b** shows the subtype organization in clusters for the second and third layers. We compare the results at the coarser level with the coarser partition into 8 broad brain regions discussed above, while the clusters at the finer level were compared with the finer organization of brain structures into 24 subregions. Different colors correspond to different brain regions, while different columns are the clusters identified by hSBM.

Looking at the histograms, we see that hSBM shows a remarkable ability in segregating different tissues solely based on the gene expression patterns of the corresponding samples. In particular, at the second hierarchical level (Figure 2**a**), hSBM turns out to be very effective in segregating cerebral gyri and lobules from the other tissues and above all in identifying the cerebellum and hippocampus samples. This resolution is further improved at the third and finer level (Figure 2**b**), where specific anatomical structures such as the cerebellar cortex, striatum, claustrum, and hypothalamus are distinctively isolated within specific clusters, while maintaining a clear distinction between cortical and non-cortical areas.

It is interesting to compare these results with those obtained using other, more standard, clustering algorithms. We used the Normalized Mutual Information (NMI)^26^ as a metric for assessing the efficacy of different algorithms in identifying the different cerebral structures (see the methods section for the definition of NMI). Notice that, since randomly shuffled sets can also yield a nonzero NMI, which increases with the number of clusters, we accounted for this effect by using a normalized version of NMI, referred to as NMI*. This normalization was achieved by dividing our NMI values by those from a basic null model that maintains the number and sizes of clusters but randomly rearranges the sample labels.

We use the NMI* score in Figure 2**c** to compare the results obtained with hSBM with the two distinct levels of samples annotations, while in Figure 2**d** we use it to evaluate the performance of different clustering algorithms, fixing the samples labeling. In the other algorithms the number of clusters is a free parameter and must be fixed by hand; we chose to fix it to achieve a number of clusters similar to that obtained with hSBM for each level of resolution.

As anticipated, the first and last layers of partition carry little information, and their score is very low. They correspond to the first and last points in the Figures 2**c** and 2**d**. Looking at Figure 2**c**, we see that the coarser ontology, referring to regions, aligns better with the second layer of hSBM clusters compared to the finer-grained labeling (subregions), while for the third layer of the hSBM clusters the order of the scores is reversed, confirming that the finer clusters correspond to substructures nested within the clusters from the previous level.

The highest value of NMI* is reached for the second layer, where, as it can be seen in Figure 2**d**, hSBM outperforms Weighted Gene Correlation Network Analysis (WGCNA), Latent Dirichlet Allocation (LDA) and hierarchical clustering.

### 2.5 Functional enrichment of topics associated to brain regions

Topics are essentially lists of genes and we may associate to each topic a functional role by performing a functional enrichment analysis. We performed functional enrichment in brain regions of interest in two steps.

- First we identified topics particularly associated to the region of interest. To this end, we used a centered version of *P*(topic | sample) defined as follows:

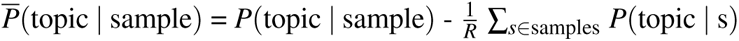

being *R* the total number of samples. In this way, we were able to compare the role of different topics in a given subset of samples and identify topics particularly enriched in regions of interest of the brain (more generally in a given subset of the samples). To allow a better visualization of this selection, the centered distributions can be represented as box plots, after grouping samples in the different subsets, i.e., different regions of interest of the brain as reported in Figure 3. The box plots presented in the first row of Figure 3 show these probability distributions for some of the 32 topics found in the second hierarchical level, which are particularly enriched in large cerebral regions. In the second and third rows we show the same enrichment patterns for the finer level of cerebral structures.
- Then we studied the functional enrichment of the genes contained in the selected topics using standard hypergeometric tests (suitably corrected for multiple testing). The results were obtained using the GSEA^27^ tool.

**Figure 3.**
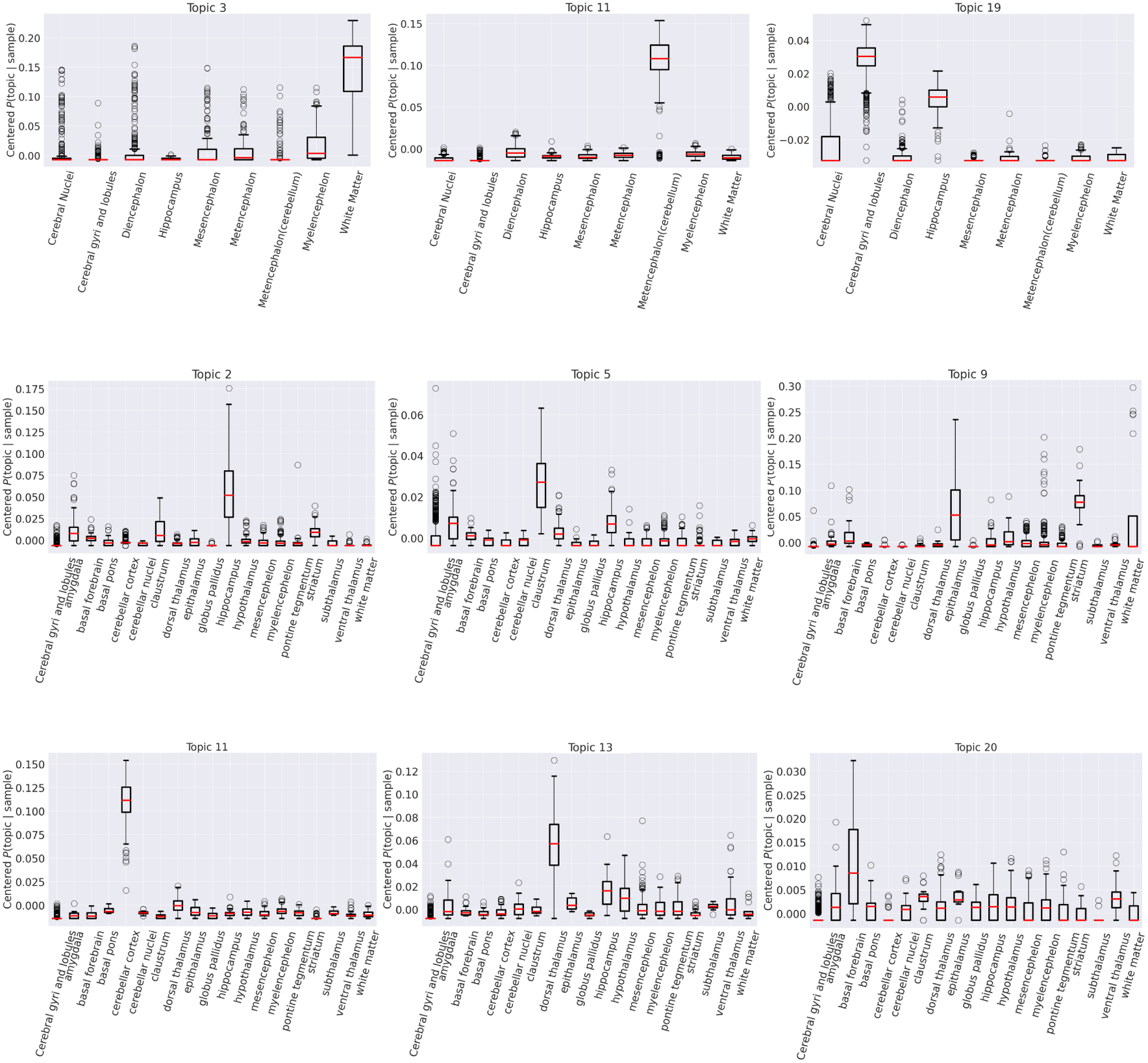
Identifying the relevant topics across samples. Distribution of *P*(topic*|*sample) for different topics from the second level of partition of hSBM across the main brain regions (box plot of the first row) and across the finer cerebral structures (box plot of the second and third rows)

Results of this analysis for the second layer, the one composed by 32 topics, are reported in Figure 3 and Table 1. Specifically Figure 3 displays in the first row a few examples of the patterns of topics enrichment obtained with the coarse ontology (the one in which the brain is divided in 8 regions), and in the second and third rows a few examples of those obtained with the finer ontology^1^.

Let us discuss in detail a few examples to better understand the whole procedure.

- Looking at the second entry of Figure 3, we see that Topic 11 is strongly enriched in the samples associated to cerebellum. Looking at the first entry of the last row we see that this enrichment is confirmed also at the level of the finer ontology. When performing functional enrichment using GSEA, we find that the 66 genes contained in this topic are indeed enriched in markers of the cerebellum with a significant p-value of 10*^−^*^8^. On one side, this can be considered as a consistency check of the whole analysis; on the other side, it suggests that the other genes within the topics, which are not annotated as cerebellum markers, are probably also involved in functions relevant for the cerebellum (see the analysis below).
- A similar results holds for Topic 2, which exhibits a probability distribution peak for the hippocampal structure (see the second row of Fig. 3) and turns out to be enriched for the GOBP MEMORY gene set, confirming the model’s results, since the hippocampus serves a critical function in memory^28^.
- Topic 9, shows enrichment for sets related to the response to endogenous stimuli and the control of movements in relation to internal or external stimuli. These sensorimotor interactions fall within the sensory processes of striatum, which, accordingly, is characterized precisely by Topic 9 genes^29^.
- Topic 19, related to the cortex both in the topic space and from the gene set enrichment analysis, is found to be associated with Alzheimer’s disease.
- The strong enrichment of Topic 5 for gene sets related to synaptic connections confirms the role of the claustrum, associated with the aforementioned topic, as a multi-modal information processing network that receives input from almost all regions of the cortex and projects back to almost all regions of the cortex^30^.

### 2.6 Comparison with the GTEx database

It is interesting to compare the topics that we identified with the brain related gene expression data collected in the Genotype-Tissue Expression (GTEx) project^31^. We report in Figure 4**a**,**b** two examples of this comparison, for two of the topics that we discussed in the previous section.

– Looking at Topic 11 results (Fig. 4**a**), it becomes clear that not only the few markers mentioned above but the entire topic is enriched in the tissues of the cerebellum and cerebellar hemisphere. It can also be appreciated from the heatmap that the genes are *specific* for these regions and show no enrichment in other regions of the brain.
– A similar pattern is also visible for Topic 19 which was associated to the cortex and that we see in Figure 4**b** to be strongly enriched in few specific region of the cortex. This makes even more interesting the connection reported in Table 1 with the Alzheimer’s disease which can be associated in this way with a specific set of GTEX samples.

**Figure 4.**
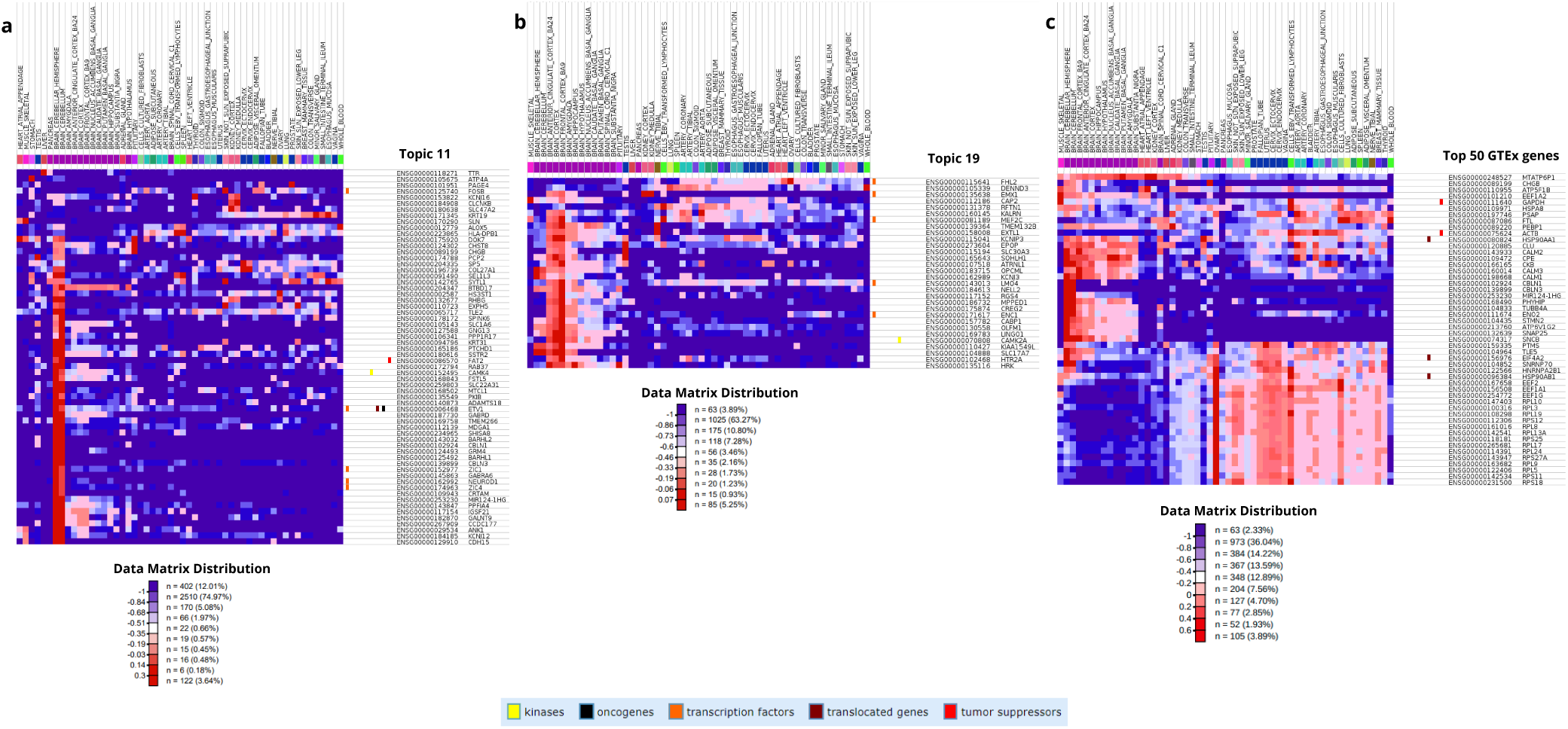
Gene expression levels of the genes in Topic 11 (**a**) and Topic 19 (**b**) (columns) from the second level of partition of hSBM output across samples of the GTEx Compendium (rows). The heatmap in **c** reports the expression profiles of the top 50 expressed genes in GTEx “Brain - Cerebellar Hemisphere” across all GTEx tissues. Each entry of the matrices is the median gene-level TPM (Transcripts Per Million) by tissue profiles from the GTEx Portal normalized with a logarithmic (log10) transformation and then mean centered. The normalized values associated with the map’s coloring and the fraction of entries for each value range are specified in the panel named “Data Matrix Distribution” for each heatmap. Genes on the rows are both identified by Ensembl Gene IDs and HGNC (HUGO) IDs. Tissue sample annotations are provided at the top matrix. The legend at the bottom includes labels that are associated with certain genes.” The heatmaps were generated using the Next-Generation Clustered Heat Maps (NG-CHM) from the Department of Bioinformatics and Computational Biology at The University of Texas MD Anderson Cancer Center^32^

Moreover, the same analysis was performed on sets of genes presented on GTEx portal as those most highly expressed in the compendium tissues. The heatmap in Figure 4**c** reports the expression profiles of the top 50 expressed genes in “Brain - Cerebellar Hemisphere”, according to the data from GTEx Compendium. It is clear that there is not such a sharp enrichment for the entire set as happened for the topic identified by hSBM, stressing that all genes identified in our topics cooperate in the regulation patterns of certain anatomical structures.

## 3 Discussion

### 3.1 hSBM is robust with respect to different gene selection choices

We used two distinct probe selection methods to evaluate the robustness of the algorithm’s clustering results with respect to the gene selection strategy. The first probe set was obtained by selecting probes exhibiting the highest variability across the merged dataset of six brains. The second selection strategy involved the independent selection of highly variable probes for each brain, subsequently merging the datasets while retaining only the probes common to all selections.

Although only 68% of the probes overlap between the two selected sets, the algorithm’s performance, in terms of similarity with the true anatomical regions of origin, remained largely consistent. To show this comparison, we computed the NMI* score for each layer of partition generated by hSBM, comparing it with the anatomical labels, as reported in the Supplementary Figure S4. For both sets, we identified four levels of resolution with similar number of clusters and similar NMI* scores, peaking at the second level where the second selection method (selecting highly variable genes separately for each brain) achieved a mean NMI* of more than 130 (i.e., 130 times those obtained from a random model).

As a last test, we also computed the NMI* score of the direct comparison of the clusters obtained with the two selection strategies (see Fig. S4). This NMI* value turned out to be very high for the second and third layers of resolution (the informative ones) and very low for the first and last layer which are essentially random partitions.

### 3.2 Comparison with the partition of Hawrylycz et al.^9^

The paper by Hawrylycz et al.^9^ serves as a reference frame in the study of AHBA data. In this paper, the authors, after a careful selection and preprocessing of the genes, identify a set of 32 “modules” using the WCGNA algorithm, which are the equivalent of our topics. It is interesting to compare their results with our topic organization. In this respect, it is impressive to notice that, with no input from our side, we find almost the same number of topics in our analysis.

We report in Supplementary Figure S5 the comparison of the gene content of the two sets of modules. Each entry in the matrix S5 reports the number of genes in common between the selected hSBM topic (on the rows) and module of ref.^9^ (on the columns). Due to the different pre-processing strategies and gene selection choices, this comparison should be taken with great care. Despite this, looking at Figure S5, a few interesting remarks can be drawn:

- In some cases, there is a mixed correspondence between topics and modules: for instance module 30 is divided into three topics (3,7,14) which are likely “submodules” of module M30. Similarly module M6 is divided into a set of topics, with a dominant role played by Topics 5, 10 and 19.
- In a few cases, there is a clear one-to-one correspondence. For instance, between Topic 11 and module M17, or between Topic 9 and module M10. These identifications show that the two partitions, even if different, are definitely not random and convey a relevant non trivial amount of information on the patterns of gene expression in the brain.
- In a completely independent way, we can use the “hub” genes proposed in^9^ as markers of the modules to map our topics into their modules. We could identify 6 of these hubs in our topics. They are reported in Table 2. A strong consistency check of the whole analysis is that the associations between topics and modules suggested by these hubs agree in almost all cases with those suggested by the overlap analysis reported in Figure S5.
- Finally, it is interesting to observe that these associations are also confirmed by the enrichment analyses reported in the previous section. For instance our Topic 11, exactly like module M17 of^9^, is enriched in the cerebellum cortex, while our Topic 13, exactly as the modulus M11 is enriched in the dorsal thalamus. Interestingly, module M6, which in^9^ is enriched in neocortex and claustrum, is associated to three topics and two of them are separately enriched in neocortex (Topic 19) and in claustrum (Topic 5) thus confirming the idea that in some cases our topics identify well defined “submodules” of the partition of^9^.

**Table 2.** Hub genes (first column) and modules (second column) according to^9^ and the corresponding topics according to our analysis (third column).

| Hub gene | Module of<sup>9</sup> | Topic |
| --- | --- | --- |
| MEF2C | M6 | 5, 12, 19 |
| NGEF | M7 | 10 |
| NTNG1 | M11 | 13 |
| SLC6A3 | M12 | 1 |
| NEFH | M15 | 26 |
| CBLN3 | M17 | 11 |

### 3.3 Mixed memberships and alternative regulatory patterns

An interesting aspect of the fuzzy membership we observe at the level of genes is that, in two cases, we have small topics which are fully included in two larger topics. In particular, Topic 30 is composed of only 8 genes, all of which also belong to Topic 19 (see Fig. S3 in the Supplementary Material). As we mentioned above, Topic 19 is associated to the neocortex. What is interesting is that *all the eight genes* are targets of the microRNA mir-142, according to the MirDIP database^33^. Mir-142 is known to be involved in neuroinflammation^34^,and specifically acts a key regulator of synaptic structure and function during neuroinflammation^35^. Accordingly, the top genes of Topic 30, i.e., those with the largest probability of being associated with this topic (see the topics content in the Supplementary Material): ‘KCNJ3’, ‘OPCML’, ‘CAMK2A’, ‘SLC17A7’, show a specific enrichment in brain functions such as “Transmission across Chemical Synapses”, “Neuroinflammation and glutamatergic signaling” or “Neurotransmitter receptors and postsynaptic signal transmission”. It is thus plausible that these genes are grouped together in Topic 30 because their expression patterns are correlated, as they are targets of mir-142.

Similarly, Topic 27, which comprises 11 genes, is fully contained in Topic 9. This Topic is enriched in basal forebrain and striatum regions (see Fig.5). Remarkably enough, also in this case all the protein-coding genes of the topic (ten out of the eleven genes of the topic^2^) are targets of a specific microRNA: mir-218 ^3^. As with the previous example, mir-218 is known to be involved in several brain functions, particularly in development and differentiation^36^.

**Figure 5.**
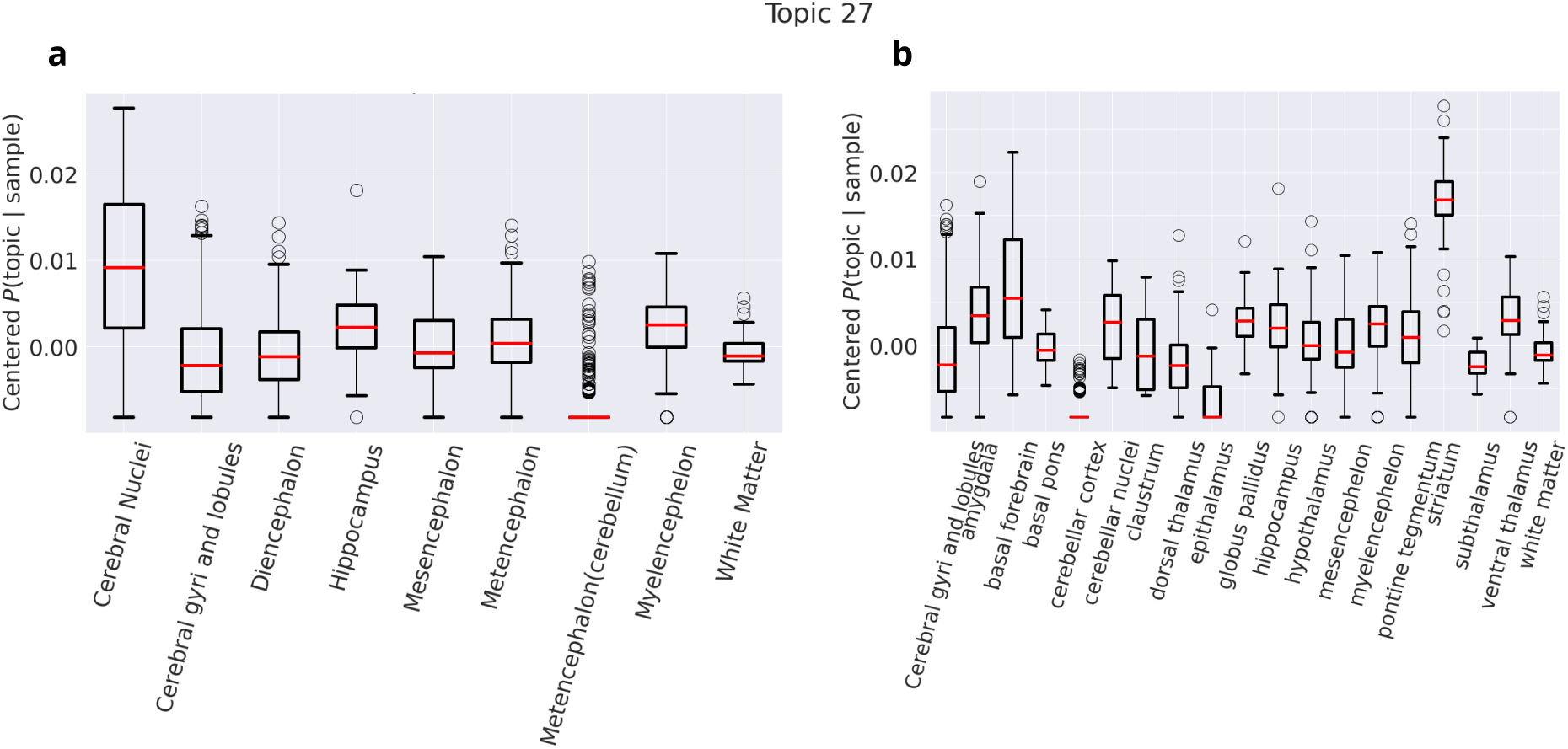
Distribution of *P*(topic*|*sample) for Topic 27. We report the probability distribution of Topic 27 from the second level of partition of hSBM, characterizing the principal brain regions (on the x-axis in **a**), and the more detailed cerebral structures in **b**.

These two examples show that the mixed membership of some of the genes may be a signature of a precise regulatory process which is active in some, but not all, regions of the brain, leading the algorithm to select it as a separate topic.

### 3.4 Mixed membership and alternative splicing: the case of MEF2C

Another interesting aspect of the mixed membership at the gene level is its possible association to alternative splicing. An interesting example is the MEF2C gene, identified as one of the “hub genes” selected by Hawrylycz et al.^9^ as representative genes of their modules. MEF2C belongs to the MEF2 (myocyte enhancer factor 2) family of Transcription Factors, consisting of four genes which play a key role in neuronal survival/apoptosis, differentiation and synaptic plasticity^37^. In particular, MEF2C has been shown to control the synapse number by inhibiting the dendritic spines growth^38^. This underscores the significant role of MEF2C in neurodevelopmental disease^39^, its specific involvement in neuropsychiatric pathologies^40^, and notably in schizophrenia^41^, as also noticed by Hawrylycz et al.^9^ when discussing the module 6, where MEF2C serves as a hub gene.

What is interesting is that MEF2C exists in the brain as different alternatively spliced isoforms^42^. The human MEF2C gene comprises 10 constitutive exons and 3 alternatively spliced exons. These alternatively spliced transcripts were proven to encode isoforms which exert distinct functions in transcriptional regulation^43^. These splicing events are finely regulated by a complex network of interactions which involve, among the others, the RNA-binding motif protein 4 (RBM4)^43^ and the alternative-splicing regulator FOX-1^42^. Accordingly, in our analysis, we observe this hub gene distributed across three different topics: Topics 5, 12 and 19, which are enriched in different brain regions and likely associated with different cell populations. This finding further highlights the flexibility of the probe-based hSBM approach.

### 3.5 hSBM is able to identify “universal” features of the dataset common to all the subjects

As mentioned in the introduction, even if the Allen Institute employed various data normalization techniques to mitigate batch effects and artefactual inter-individual differences, a significant level of intrinsic donor-specific variation persists^8^ and represents a major obstacle in the search for *universal* features common to all subjects.

Some studies have addressed these differences, mainly evident in cortical tissues, by applying subject-specific normalization prior to data aggregation (^44^,^45^,^46^,^47^), while others have compared various normalization methods after aggregating donor data (^48^,^8^). In this study, we chose a different strategy, looking for universal patterns in gene expression data without a previous subject-specific normalization. We leveraged the probabilistic nature of topic modeling, specifically the flat prior version represented by hSBM.

To evaluate the ability of different algorithms to identify universal features in the dataset, we assessed whether it was possible to discriminate individual donor brains by projecting tissue samples from all six brains into a two-dimensional transcriptional principal components space. As mentioned before, the output of some of the algorithms includes a transformation of samples from the gene space to the topic space, through the probability assigned to a sample of being driven by the cooperative action of a specific set of genes. Thus, we can then consider new features for each sample, that is the probabilities of interacting with the found topics, and assign them a position in the new multidimensional topic space. Figures 6 **a,b,c** show the result of the principal component analysis (PCA) on the topic space, plotting the values of the first two principal components for each tissue sample from all six donor brains.

**Figure 6.**
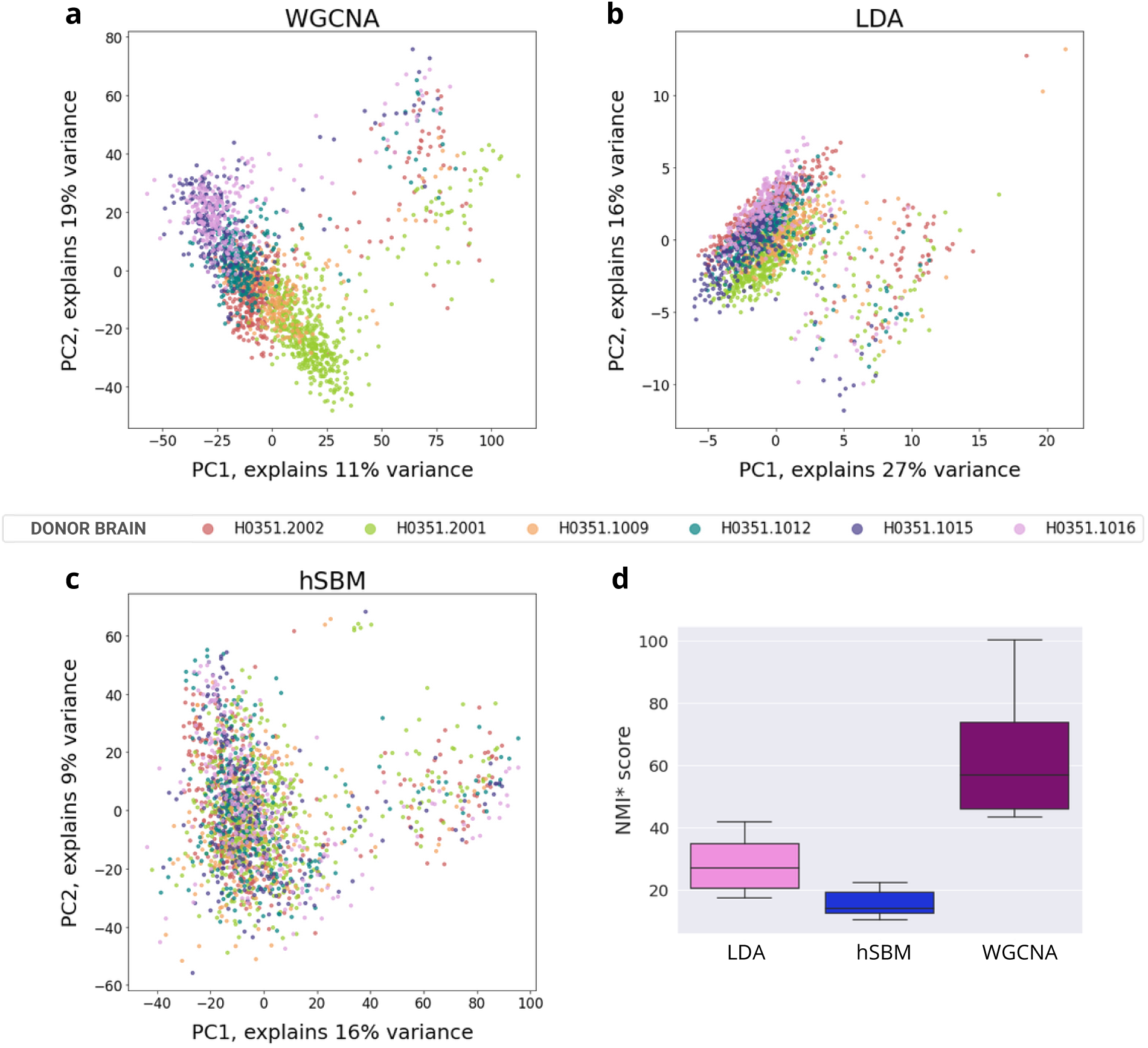
(**a**,**b**,**c**)Features, obtained as output after performing Weighted Gene Correlation Network Analysis (WGCNA) (**a**), Latent Dirichlet Allocation (LDA) (**b**) and Hierarchical Stochastic Block Model (hSBM) (**c**) on gene expression data, in principal component (PC) space. Data from different donors are represented in different colors. In panel **a** and **b** samples from different subjects occupy different parts of the low-dimensional topic space. Panel **c** shows that samples in hSBM results no longer segregate by donor. The boxplot in **d** presents the NMI* scores (normalized using a null model), calculated from ten realizations of each algorithm at a level of partitioning of samples into clusters of the same order of magnitude as the number of donors. These scores compare the clustering results of the algorithms with the sample membership to the six donor brains. The highest similarity between the two partitions is observed in the outputs of WGCNA, while the clustering produced by hSBM shows only minimal resemblance to the division based on the six brains, corroborating the findings from the PCA analysis.

By applying this analysis to three different algorithms, it was observed that the unsupervised projection of samples into the topic space reveals the underlying dimensions of variance among all samples. For the WGCNA and LDA outcomes (Fig. 6 **a** and **b**), this approach effectively distinguishes the six donors, regardless of the tissue location in the brain, indicating that these methods strongly focus on inter-individual differences, especially for cortical samples.

In contrast, the dimensionality reduction of the topic space of hSBM (Fig. 6 **c**) shows that samples no longer segregate by donor.

A quantitative assessment of this observation is presented in Figure 6 **d**, where we report the NMI* scores derived from comparing the clustering at the second hierarchical level (which corresponds to a number of sample clusters comparable to the number of patients) for each algorithm. This comparison is made against the sample labeling based on the six donor brains. It is clear that the similarity estimate between the two partitions is notably higher for the clustering performed by WGCNA compared to that obtained with hSBM. This comparison supports the conclusions drawn from the dimensionality reduction in the topic space previously illustrated, suggesting that hSBM may integrate these inter-individual differences differently or focus less on them, potentially capturing more universal patterns across the dataset.

## 4 Methods

### The AHBA dataset

The AHBA microarray gene expression data includes 3,702 samples from six neurotypical adult brains. Samples were obtained from cortical, subcortical, brainstem and cerebellar regions in each brain and profiled for genome-wide gene expression using custom microarrays containing a total of 58 692 probes. The AHBA provides an annotation table to map probes to genes. In the original release of the AHBA dataset 48 171 probes out of the 58 692 were associated to a gene, leading to a set of 20,787 unique genes with expression measurements. were associated to a gene, resulting in a set of 20 787 unique genes with expression measures, of which, 93% are associated to more than one probe. As we shall see below, after preprocessing and reannotation these numbers were sligthly reduced.

Besides gene expression data, the AHBA assigns a binary indicator to each probe in each sample to determine whether it detects an expression signal above background noise. This information plays an important role in the preprocessing pipeline. Each tissue sample is associated with a unique numerical structure ID, a descriptive name, and a structural label. Since Since tissue samples were not collected in a spatially uniform manner across all donor brains, each brain may contribute differing quantities of samples to any specific brain region. This makes the goal of finding common gene expression patterns across different brains even more challenging. Further details on the AHBA can be found in the section S1 of the Supplementary Material.

### Data Preprocess and Gene Filtering

The AHBA contains over 20 000 unique genes, but only a small portion is expected to exhibit consistent regional variations in expression throughout the brain. Before choosing a filtering technique, we decided to follow the first two steps of the Arnatkeviciute et al.’s pipeline^8^, which involve verifying the probe-gene annotation and filtering out probes that do not exceed background noise.

Microarray experiments utilize probe sequences representing unique DNA segments, allocated to genes based on genome sequencing databases. Although platforms like AHBA offer annotation tables mapping probes to genes, as suggested by Arnatkeviciute et al., these data become outdated with sequencing database updates and an accurate probe-to-gene mapping is crucial for biologically meaningful results. Therefore, a re-assignment of probes using the latest information was necessary. This was performed through the Re-annotator software, which uses probe sequences and mRNA reference database information to identify genes that match specific microarray probe sequences^8^.

We found that, out of a total of 58 692 probe sequences, 45 907 (78%) are uniquely associated with a gene and can be linked to an entrez ID, which is a unique identifier generated by the Entrez Gene database at the National Center for Biotechnology Information (NCBI). Approximately 19% of the probes were not associated with any gene, while 3% are mapped to multiple genes, making it impossible to make a single choice. These were eliminated from the subsequent analysis.

At the next step, the aforementioned AHBA binary indicator [intensity based filtering (IBF)] was used to remove a fixed percentage of probes with lowest signal intensities. This is because microarray experiments often encounter background noise due to non-specific hybridization, and where signal levels are very close to the background noise (lower hybridization intensities), the variability in measured intensity values increases^49^. We exclude probes that did not exceed the background in at least 25% of all samples across all subjects, thus retaining 75% of the probes, which amounts to 34 522 (corresponding to 16 658 genes) probes out of 45 907. One may fear a loss of information when applying this filter, however it was shown in^50^, that the genes which are eliminated by IBF are typically associated with general cellular, immunological, and metabolic processes, which are not brain-specific.

After performing re-annotation and IBF, 73% of the remaining genes are mapped to at least two probes. It happens that different probes associated with a gene, contrary to what one might expect, exhibit diverse gene expression patterns across samples, as demonstrated by Arnatkeviciute et al. This makes it difficult to assign a precise expression value to a given gene. To avoid this problem, we decided, as mentioned above, to use the single probes as entries for our algorithm.

However, a further filtering step on the probe side was necessary in order to reduce the computational cost of the algorithms. To accomplish this, we opted for selecting probes that exhibit highest variability, being differentially expressed among samples, as is usually done for gene filtering. A major consequence of this filter is to eliminate *”housekeeping probes”*, i.e. probes linked to genes that are actively transcribed and translated to a relative high level in all cell and tissues. Generally, they encode proteins and enzymes that are essential for cell life and are not very useful to identify the cell population or tissue to which the sample belongs. To make a comparison with text mining problem, housekeeping genes have the same role as conjunctions, they are present in every text but do not in any way identify documents topic.

Given the particular nature of our dataset (six different brains merged together) we have several different ways to apply the highly variable filter. As we mentioned in the results section, we chose two orthogonal strategies to performed this selection. The first choice was to select probes with the highest variability within the merged dataset of the six brains. The second strategy was to independently select highly variable probes for each brain, and merge the datasets by keeping only the probes common to all selections. In both cases we selected the 4 000 most variable probes. In the first case the highly variable probes are mapped to 1 210 different genes, while in the second case they are associated to 1 743 genes. The overlap between the two selections turns out to be of only 2 750 probes corresponding to 1 001 genes.

The two choices are “orthogonal” in the sense that the first has the effect of keeping as low as possible the inter-individual differences among the six brains, while the second has the effect of magnifying these differences. This “orthogonality” is clearly visible if we look at the overlap between the two selections which turns out to be of only 2 750 probes (out of the 4000 of the two sets) corresponding to 1 001 genes. The comparison between the two choices (details of which are reported in the section S2 of the supplementary material) allows us to test the robustness of our results.

### Hierarchical Stochastic Block Model

We adjust the hierarchical Stochastic Block Model to suit the analysis of gene expression data from brain tissue. The code to run hierarchical stochastic block model on a bipartite network is available at the repository: https://github.com/fvalle1/hSBM Topicmodel/tree/develop.

Hierarchical stochastic block model (hSBM) is a kind of generative model aimed at maximizing the probability that the model *θ* accurately describe the data *A*

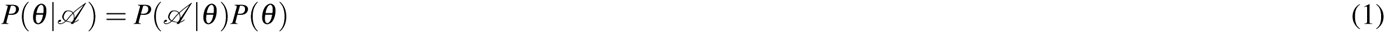

using a non-parametric approach. In our case *A* is the gene expression matrix and the entries *A_i j_* represents the expression level of gene *i* in the sample *j*. It is easy to see that the *A* can be interpreted as the adjacency matrix of a bipartite network composed by genes on one side and samples on the other side (see Fig.1). The edges of this network are weighted by the gene expression levels.

The goal of the algorithm is to minimize the description length Σ = *−lnP*(*A |θ*) *− lnP*(*θ*) of the model. We set the algorithm so as to minimise the description length Σ repeatedly and chose the model that resulted in the shortest description length.

The model outputs the probability distributions *P*(topic*|*sample) and *P*(probe*|*topic). These probabilities are defined in terms of the program’s entries as follows:

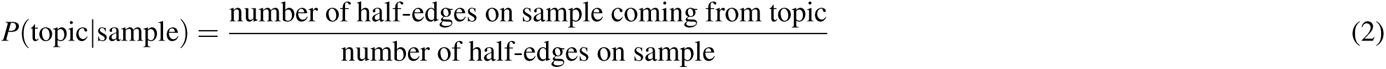

and

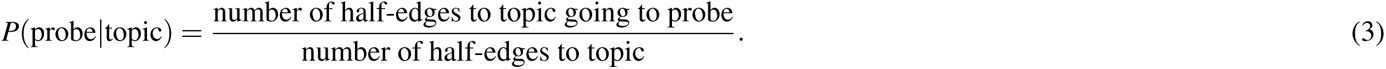

Considering the probability of the probe given the topic, a probability of the gene given the topic has been defined by selecting, among all possible probes associated with a gene that belongs to the same topic, the highest value. However, it is clear that probes associated with the same gene can also be divided into different topics, thus causing a gene to appear in multiple topics.

### Normalised Mutual Information (*NMI*)

We used the Normalised Mutual Information *NMI*^51^ to assess the performance of different clustering strategies^52^. Given a collection *C* of labeled samples and a partition *K* dividing these samples into clusters, the NMI is calculated as the harmonic average of homogeneity *h* and completeness *C* :

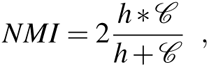

where the homogeneity is defined as

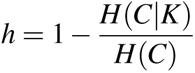

and the completeness as

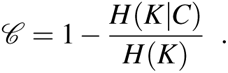

In these definitions *H*(*C*) and *H*(*K*) are the usual Shannon entropies associated to the partitions *C* and *K*; *H*(*C|K*) and *H*(*K|C*) are defined as:

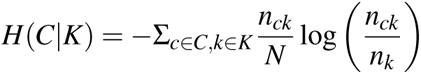

and

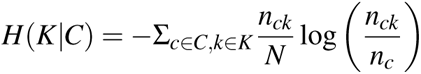

respectively, where *n_c_*is the number of samples of the region *c*, *n_k_*the number of samples in the cluster *k* and *n_ck_*the number of samples of the region *c* in the cluster *k*.

Since randomly shuffled sets can also yield a nonzero NMI, which increases with the number of clusters, we accounted for this effect by using a normalized version of NMI, referred to as NMI*. This normalization was achieved by dividing our NMI values by those from a basic null model that maintains the number and sizes of clusters but randomly rearranges the sample labels.

### Comparison with the GTEX database

The comparison with the GTEX data was performed using a GSEA portal function which allows the production of heatmaps from a user-provided gene list against the samples of several compendia of expression data and in particular with those of the Genotype-Tissue Expression (GTEx) project^31^ discussed in the text. This is done through the Next-Generation Clustered Heat Maps (NG-CHM) tool from the Department of Bioinformatics and Computational Biology at The University of Texas MD Anderson Cancer Center^32^. The data used for the anlysis described in the manuscript were obtained from the GTEx portal https://gtexportal.org/home/license on 23/05/2024.

### Implementation of LDA, WCGNA and hierarchical algorithms

In this study, we also employed alternative clustering techniques to compare their performance with that of hSBM. For each method, we aimed to select hyperparameters that would yield a number of clusters comparable to those generated by hSBM, which is completely non-parametric.

Weighted Gene Correlation Network Analysis (WGCNA) was run using the dedicated R package available at https://cran.r-project.org/web/packages/WGCNA/index.html. The analysis was conducted with default parameters: the *power* was set to the minimum value at which the scale-free topology fit index curve plateaued, *minModuleSize* was set to 5, and *mergeCutHeight* to 0.2. WGCNA organizes genes into *modules*, which we regarded as topics. To derive clusters, we pruned the tree constructed from these modules to estimate sample distances.

Latent Dirichlet Allocation (LDA)^17^ was performed using the implementation provided *sklearn*^53^. The model was set up using the default configuration for the parameters *α* and *β*, which respectively represent the Dirichlet distribution parameters used to sample words (for a topic) and topics (for a document). Both parameters were set to 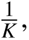 where *K* is the number of topics. The value of *K* was determined from the number of clusters produced as hSBM output. When managing LDA output, we chose the *argmax* of *P*(topic *|* sample) to determine cluster assignments. Therefore, since LDA does not have a hierarchical output, it was run a number of times equal to the number of levels returned in output by hSBM, each time setting as the parameter of LDA a number of clusters of the same order of magnitude as those from hSBM. This allowed for a level-by-level comparison of the algorithms’ performance.

Hierarchical clustering was carried out with *sklearn*, employing the Euclidean metric and complete linkage. We adjusted the clustering to align with the number of clusters determined by the hSBM. Unfortunately, this approach did not yield any information about the genes side.

### Results visualization

All the plots reported in Figures 2,3,5 were created using the Python package *topicpy*, implemented for topic modelling pre and post processing and available at https://pypi.org/project/topicpy/.

## 5 Conclusions

In this paper we presented a new approach to the study of brain gene expression data based on a hierarchical version of Stochastic Block Modeling. We tested it on a dataset of the Allen Human Brain Atlas and compared its performances with that of other data-mining tools, such as WCGNA, previously applied to the same dataset. The main goal of our analysis was to show that a careful choice of priors, implicit in different models, allows to identify universal features in the dataset, and to overcome the problem of inter-individual differences, which is a major obstacle in analysing gene expression data of complex tissues like those composing the central nervous system. It would be interesting to extend this strategy to the much larger datasets which are now available^1–4^. The major bottleneck is the computational cost of hSBM, which is definitely larger than that of the other competing algorithms we tested. However, there are two positive considerations: first, most of the recently available data are single-cell data which are very sparse. In this situation, it can be shown that the computational cost of hSBM grows only as *O*(*VLn*^2^*V*) (where *V* is the total number of vertices of the bipartite network, i.e. the number of samples plus the number of genes)^24^ and in fact hSBM was recently successfully applied on a large database of cancer single-cell data^20^. The second is that topic modeling, and in particular SBM-based models, is a very active line of research, and remarkable improvements in the performance of this class of algorithms have been obtained using for instance merge-split Markov Chains^54^, emerged consensus over SBM approach^55^.

## Acknowledgements

We would like to acknowledge Pietro Liò and Loredana Martignetti for useful discussions.

## Author contributions statement

M.C. and M.O. conceived and supervised the study; L.P. and F.V. identified key results and developed the methodologies; L.P. analyzed the data; The first draft of the manuscript was written by M.C. and L.P. and all authors commented on previous versions of the manuscript. All authors read and approved the final manuscript.

## Footnotes

1 For a better visualization of the results, in line with the reference atlases of AHBA, we merged the labels corresponding to the parietal lobe, frontal lobe, temporal lobe, occipital lobe, limbic lobe and insular lobe in a single entry named ‘Cerebral gyri and lobules’ thus reducing the original 24 subregions to 19 entries in Figure 3.

2 The only missing target in the topic is the long non coding gene LINC0982 which is missing because the mirDIP database does not contain lncRNA targets.

3 More precisely by the two components of the mir-218 family: mir-218a and mir218b which have the same seed sequence and are thus considered as a single mature miRNA by mirDIP

## References

1. Callaway, E. M. et al. A multimodal cell census and atlas of the mammalian primary motor cortex. Nature 598, 86–102 (2021). URL 10.1038/s41586-021-03950-0.

2. Luo, C. et al. Single nucleus multi-omics identifies human cortical cell regulatory genome diversity. Cell Genomics 2, 100107 (2022). URL https://www.sciencedirect.com/science/article/pii/S2666979X22000271.

3. Emani, P. S. et al. Single-cell genomics and regulatory networks for 388 human brains. Science 384 (2024). URL 10.1126/science.adi5199.

4. Huuki-Myers, L. A. et al. A data-driven single-cell and spatial transcriptomic map of the human prefrontal cortex. Science 384 (2024). URL 10.1126/science.adh1938.

5. Keil, J. M., Qalieh, A. & Kwan, K. Y. Brain transcriptome databases: A user’s guide. The Journal of Neuroscience 38, 2399–2412 (2018). URL 10.1523/JNEUROSCI.1930-17.2018.

6. Hawrylycz, M. J. et al. An anatomically comprehensive atlas of the adult human brain transcriptome. Nature 489, 391–399 (2012). URL 10.1038/nature11405.

7. Shen, E. H., Overly, C. C. & Jones, A. R. The allen human brain atlas. Trends in Neurosciences 35, 711–714 (2012). URL 10.1016/j.tins.2012.09.005.

8. Arnatkeviciūtė, A., Fulcher, B. D. & Fornito, A. A practical guide to linking brain-wide gene expression and neuroimaging data. NeuroImage 189, 353–367 (2019). URL 10.1016/j.neuroimage.2019.01.011.

9. Hawrylycz, M. et al. Canonical genetic signatures of the adult human brain. Nature Neuroscience 18, 1832–1844 (2015). URL 10.1038/nn.4171.

10. Richiardi, J. et al. Correlated gene expression supports synchronous activity in brain networks. Science 348, 1241– 1244 (2015). URL https://www.science.org/doi/abs/10.1126/science.1255905. https://www.science.org/doi/pdf/10.1126/science.1255905.

11. Cioli, C., Abdi, H., Beaton, D., Burnod, Y. & Mesmoudi, S. Differences in human cortical gene expression match the temporal properties of large-scale functional networks. PLoS ONE 9, e115913 (2014). URL 10.1371/journal.pone.0115913.

12. Eising, E. et al. Gene co-expression analysis identifies brain regions and cell types involved in migraine pathophysiology: a gwas-based study using the allen human brain atlas. Human Genetics 135, 425–439 (2016). URL 10.1007/s00439-016-1638-x.

13. Kumar, A. et al. Age-associated changes in gene expression in human brain and isolated neurons. Neurobiology of Aging 34, 1199–1209 (2013). URL 10.1016/j.neurobiolaging.2012.10.021.

14. Trabzuni, D. et al. Widespread sex differences in gene expression and splicing in the adult human brain. Nature Communications 4 (2013). URL 10.1038/ncomms3771.

15. Gerlach, M., Peixoto, T. P. & Altmann, E. G. A network approach to topic models. Science Advances 4 (2018). URL 10.1126/sciadv.aaq1360.

16. Valle, F., Osella, M. & Caselle, M. A topic modeling analysis of tcga breast and lung cancer transcriptomic data. Cancers 12, 3799 (2020). URL 10.3390/cancers12123799.

17. Blei, D. M., Ng, A. Y. & Jordan, M. I. Latent dirichlet allocation. J. Mach. Learn. Res. 3, 993–1022 (2003). URL https://dl.acm.org/doi/10.5555/944919.944937.

18. Liu, L., Tang, L., Dong, W., Yao, S. & Zhou, W. An overview of topic modeling and its current applications in bioinformatics. SpringerPlus 5 (2016). URL 10.1186/s40064-016-3252-8.

19. Valle, F., Osella, M. & Caselle, M. Multiomics topic modeling for breast cancer classification. Cancers 14, 1150 (2022). URL 10.3390/cancers14051150.

20. Malagoli, G., Valle, F., Barillot, E., Caselle, M. & Martignetti, L. Identification of interpretable clusters and associated signatures in breast cancer single-cell data: A topic modeling approach. Cancers 16, 1350 (2024). URL 10.3390/cancers16071350.

21. Lazzardi, S. et al. Emergent statistical laws in single-cell transcriptomic data. Physical Review E 107 (2023). URL 10.1103/PhysRevE.107.044403.

22. Fortunato, S. & Hric, D. Community detection in networks: A user guide. Physics Reports 659, 1–44 (2016). URL 10.1016/j.physrep.2016.09.002.

23. Langfelder, P. & Horvath, S. Wgcna: an r package for weighted correlation network analysis. BMC Bioinformatics 9 (2008). URL 10.1186/1471-2105-9-559.

24. Peixoto, T. P. Hierarchical block structures and high-resolution model selection in large networks. Physical Review X 4 (2014). URL 10.1103/PhysRevX.4.011047.

25. Ward, J. H. Hierarchical grouping to optimize an objective function. Journal of the American Statistical Association 58, 236–244 (1963). URL 10.1080/01621459.1963.10500845.

26. Shi, H., Gerlach, M., Diersen, I., Downey, D. & Amaral, L. A new evaluation framework for topic modeling algorithms based on synthetic corpora. In Chaudhuri, K. & Sugiyama, M. (eds.) Proceedings of the Twenty-Second International Conference on Artificial Intelligence and Statistics, vol. 89 of Proceedings of Machine Learning Research, 816–826 (PMLR, 2019). URL https://proceedings.mlr.press/v89/shi19a.html.

27. Subramanian, A. et al. Gene set enrichment analysis: A knowledge-based approach for interpreting genome-wide expression profiles. Proceedings of the National Academy of Sciences 102, 15545–15550 (2005). URL https://www.pnas.org/doi/abs/10.1073/pnas.0506580102. https://www.pnas.org/doi/pdf/10. 1073/pnas.0506580102.

28. Lisman, J. et al. Viewpoints: how the hippocampus contributes to memory, navigation and cognition. Nature Neuroscience 20, 1434–1447 (2017). URL 10.1038/nn.4661.

29. de la Torre-Martinez, R., Ketzef, M. & Silberberg, G. Ongoing movement controls sensory integration in the dorsolateral striatum. Nature Communications 14 (2023). URL 10.1038/s41467-023-36648-0.

30. Nikolenko, V. N. et al. The mystery of claustral neural circuits and recent updates on its role in neurodegenerative pathology. Behavioral and Brain Functions 17 (2021). URL 10.1186/s12993-021-00181-1.

31. Lonsdale, J. et al. The genotype-tissue expression (gtex) project. Nature Genetics 45, 580–585 (2013). URL 10.1038/ng.2653.

32. Ryan, M. C. et al. Interactive clustered heat map builder: An easy web-based tool for creating sophisticated clustered heat maps. F1000Research 8, 1750 (2019). URL 10.12688/f1000research.20590.1.

33. Tokar, T. et al. mirdip 4.1—integrative database of human microrna target predictions. Nucleic Acids Research 46, D360–D370 (2017). URL 10.1093/nar/gkx1144.

34. Korotkov, A. et al. Increased expression of mir142 and mir155 in glial and immune cells after traumatic brain injury may contribute to neuroinflammation via astrocyte activation. Brain Pathology 30, 897–912 (2020). URL 10.1111/bpa.12865.

35. Mandolesi, G. et al. mir-142-3p is a key regulator of il-1-dependent synaptopathy in neuroinflammation. The Journal of Neuroscience 37, 546–561 (2016). URL 10.1523/JNEUROSCI.0851-16.2016.

36. Sempere, L. F. et al. Expression profiling of mammalian micrornas uncovers a subset of brain-expressed micrornas with possible roles in murine and human neuronal differentiation. Genome Biology 5 (2004). URL 10.1186/gb-2004-5-3-r13.

37. Allaway, K. C. et al. Genetic and epigenetic coordination of cortical interneuron development. Nature 597, 693–697 (2021). URL 10.1038/s41586-021-03933-1.

38. Paciorkowski, A. R. et al. Mef2c haploinsufficiency features consistent hyperkinesis, variable epilepsy, and has a role in dorsal and ventral neuronal developmental pathways. neurogenetics 14, 99–111 (2013). URL 10.1007/s10048-013-0356-y.

39. Cosgrove, D. et al. Genes influenced by mef2c contribute to neurodevelopmental disease via gene expression changes that affect multiple types of cortical excitatory neurons. Human Molecular Genetics 30, 961–970 (2020). URL 10.1093/hmg/ddaa213.

40. Zhang, Z. & Zhao, Y. Progress on the roles of mef2c in neuropsychiatric diseases. Molecular Brain 15 (2022). URL 10.1186/s13041-021-00892-6.

41. Mitchell, A. C. et al. Mef2c transcription factor is associated with the genetic and epigenetic risk architecture of schizophrenia and improves cognition in mice. Molecular Psychiatry 23, 123–132 (2017). URL 10.1038/mp.2016.254.

42. Hakim, N. H. A., Kounishi, T., Alam, A. H. M. K., Tsukahara, T. & Suzuki, H. Alternative splicing ofmef2cpromoted by fox-1 during neural differentiation in p19 cells. Genes to Cells 15, 255–267 (2010). URL 10.1111/j.1365-2443.2009.01378.x.

43. Lin, J.-C. Rbm4-mef2c network constitutes a feed-forward circuit that facilitates the differentiation of brown adipocytes. RNA Biology 12, 208–220 (2015). URL 10.1080/15476286.2015.1017213.

44. Liu, Z. et al. The functional and genetic associations of neuroimaging data: a toolbox. BioRxiv (2017). URL 10.1101/178640.

45. Romero-Garcia, R. et al. Structural covariance networks are coupled to expression of genes enriched in supragranular layers of the human cortex. NeuroImage 171, 256–267 (2018). URL 10.1016/j.neuroimage.2017.12.060.

46. Burt, J. B. et al. Hierarchy of transcriptomic specialization across human cortex captured by structural neuroimaging topography. Nature Neuroscience 21, 1251–1259 (2018). URL 10.1038/s41593-018-0195-0.

47. McColgan, P. et al. Brain regions showing white matter loss in huntington’s disease are enriched for synaptic and metabolic genes. Biological Psychiatry 83, 456–465 (2018). URL 10.1016/j.biopsych.2017.10.019.

48. Parkes, L., Fulcher, B. D., Yücel, M. & Fornito, A. Transcriptional signatures of connectomic subregions of the human striatum. *Genes*, Brain and Behavior 16, 647–663 (2017). URL 10.1111/gbb.12386.

49. Quackenbush, J. Microarray data normalization and transformation. Nature Genetics 32, 496–501 (2002). URL 10.1038/ng1032.

50. Gillis, J., Mistry, M. & Pavlidis, P. Gene function analysis in complex data sets using erminej. Nature Protocols 5, 1148–1159 (2010). URL 10.1038/nprot.2010.78.

51. Rosenberg, A. & Hirschberg, J. V-measure: A conditional entropy-based external cluster evaluation measure. In Conference on Empirical Methods in Natural Language Processing (2007). URL https://api.semanticscholar.org/ CorpusID:14153811.

52. Shi, H., Gerlach, M., Diersen, I., Downey, D. & Amaral, L. A new evaluation framework for topic modeling algorithms based on synthetic corpora. In Chaudhuri, K. & Sugiyama, M. (eds.) Proceedings of the Twenty-Second International Conference on Artificial Intelligence and Statistics, vol. 89 of Proceedings of Machine Learning Research, 816–826 (PMLR, 2019). URL https://proceedings.mlr.press/v89/shi19a.html.

53. Hoffman, M. D., Blei, D. M. & Bach, F. Online learning for latent dirichlet allocation. In Proceedings of the 23rd International Conference on Neural Information Processing Systems - Volume 1, NIPS’10, 856–864 (Curran Associates Inc., Red Hook, NY, USA, 2010).

54. Peixoto, T. P. Merge-split markov chain monte carlo for community detection. Physical Review E 102 (2020). URL 10.1103/PhysRevE.102.012305.

55. Peixoto, T. P. Revealing Consensus and Dissensus between Network Partitions. Physical Review X 11, 021003 (2021). URL https://link.aps.org/doi/10.1103/PhysRevX.11.021003.

