## Supplementary Material for "Topic Modeling analysis of the Allen Human Brain Atlas"

#### S1 Details on the AHBA dataset

The AHBA microarray gene expression data consists of 3702 samples from six neurotypical adult brains. Several hundred samples (mean ± standard deviation: 617 ± 241) were collected from cortical, subcortical, brainstem and cerebellar regions in each brain and profiled for genome-wide gene expression using custom Agilent 8 × 60K cDNA chip which consists of a standard Whole Human Genome Microarray Kit, 4 × 44K (Design ID: 014850) and more than 18,000 custom-generated probes created specifically for AHBA in order to increase the genetic coverage. The AHBA provides the annotation table to map probes to genes. In the original release of the AHBA 48'171 probes out of a total of 58'692 are associated to a gene, resulting in a set of 20'787 unique genes with expression measures, of which, 93% were associated with more than one probe, since a single gene's expression levels can be assessed using various probes, each corresponding to different segments of the gene sequence.
As we mentioned in the main text, after preprocesing and reannotation only 45,907 (78%) of the probes were uniquely associated with a gene and could be linked to an entrez ID, In addition to gene expression data, the AHBA assigns a binary indicator to each probe in each sample to determine whether it detects an expression signal above background noise. This assignment is done on the basis of two criteria: 1) a two-sided t-statistic comparing the mean signal of a probe to the background ($p < 0.01$); and 2) retaining only background-subtracted signals that are 2.6 × above the standard deviation of the background.
Each probe in the AHBA is associated with a numerical ID and a platform-specific label or name.
As mentioned in the main text probe-level data are accessible for each of the 3702 tissue samples collected from various regions of the brain. Different brain regions were sampled across each of the six AHBA donors to maximize spatial coverage. Each tissue sample is associated with a unique numerical structure ID, a descriptive name, and a structural label. Moreover, it contains MRI voxel coordinates in the native image space and MNI coordinates in a standardized space, simplifying the process of aligning samples with the stereotaxic space. Spatial distribution of tissue samples across individual brains in the AHBA is not uniform. Therefore, various brains may contribute differing quantities of samples to any specific brain region (see Figure S1)
The AHBA also provides RNA-seq data for a smaller selection of samples from two of the previous donors (120 samples each) which were not used in the present work. This type of data, available for over 22,000 genes, are available both in fragment count (representing the number of reads corresponding to a specific gene) and TMP (Transcripts Per Kilobase Million - normalized read count adjusted for read and transcript length) formats,
Deatils on the six donors are reported in the Table S1.

| **Donor** | **Age** | **Sex** | **Race/Ethnicity** | **Medical Condition** | **Postmortem interval**[b] |
|---|---|---|---|---|---|
| H0351_1009 | 57 | Male | Caucasian | History of atherosclerotic cardiovascular disease. | 25.5 hours |
| H0351_1012 | 31 | Male | Caucasian | Sudden cardiac arrest. Benign spindle cell proliferation and dystrophic calcification in temporal horn of lateral ventricle. | 17.5 hours |
| H0351_1015 | 49 | Female | Hispanic | Splenectomy, hypothyroidism treated with Levothroid; modest numbers of hemosiderin laden macrophages noted in Virchow-Robin spaces in parietal and occipital lobes, mild arteriosclerosis. | 30 hours |
| H0351_1016 | 55 | Male | Caucasian | Coronary artery atherosclerosis, prescriptions for clotting and high cholesterol. | 18 hours |
| H0351_2001[a] | 24 | Male | African American | History of asthma | 23 hours |
| H0351_2002[a] | 39 | Male | African American | Possible small pituitary adenoma Microneuropathology; single neurofibrillary tangle in entorhinal cortex. | 10 hours |

Table S1: **Profiles of the six adult donors in AHBA.** For more detailed information about the six subjects, screening tests and quality control measures performed to ensure the tissue and RNA met quality control criteria see *Case Qualification and Donor Profiles in the Allen Human Brain Atlas.*

[a]These donors have tissue samples collected across both left and right hemispheres, different from other donors that have samples only within the left hemispheres.
[b] Postmortem interval is defined as the time period from the time of death to the time the tissue is frozen.

| Brain Region (coarse-grained ontology) | | Brain Subregion (fine-grained ontology) | Brain H0351.2002 # of samples | Brain H0351.2001 # of samples | Brain H0351.1009 # of samples | Brain H0351.1012 # of samples | Brain H0351.1015 # of samples | Brain H0351.1016 # of samples | Total # of samples |
|---|---|---|---|---|---|---|---|---|---|
| Cerebral gyri and lobules | | Frontal Lobe | 121 | 165 | 46 | 69 | 64 | 64 | 529 |
| | | Parietal Lobe | 70 | 77 | 39 | 43 | 30 | 29 | 288 |
| | | Occipital Lobe | 43 | 34 | 24 | 39 | 37 | 43 | 220 |
| | | Temporal Lobe | 87 | 157 | 37 | 72 | 51 | 67 | 471 |
| | | Limbic Lobe | 43 | 36 | 30 | 34 | 35 | 31 | 209 |
| | | Insular Lobe | 6 | 12 | 4 | 8 | 7 | 6 | 43 |
| | | Hippocampus | 54 | 61 | 14 | 22 | 17 | 20 | 188 |
| Cerebral Nuclei | | Striatum | 46 | 47 | 16 | 24 | 18 | 18 | 169 |
| | | Globus Pallidus | 14 | 11 | 4 | 2 | 3 | 5 | 39 |
| | | Basal Forebrain | 10 | 7 | 4 | 9 | 6 | 11 | 47 |
| | | Amygdala | 24 | 16 | 11 | 6 | 7 | 10 | 74 |
| | | Claustrum | 11 | 17 | 5 | 7 | 5 | 2 | 47 |
| Diencephalon | | Dorsal Thalamus | 44 | 47 | 21 | 23 | 16 | 20 | 171 |
| | | Ventral Thalamus | 13 | 7 | 6 | 5 | 5 | 1 | 37 |
| | | Hypothalamus | 22 | 9 | 19 | 20 | 16 | 19 | 105 |
| | | Subthalamus | 3 | 3 | 2 | 2 | 1 | 3 | 14 |
| | | Epithalamus | 2 | 8 | 2 | 2 | 0 | 1 | 15 |
| | | Mesencephelon | 61 | 48 | 8 | 22 | 24 | 18 | 181 |
| Metence-phalon | Cerebel-lum | Cerebellar Cortex | 76 | 41 | 40 | 44 | 60 | 76 | 337 |
| | | Cerebellar Nuclei | 7 | 12 | 2 | 4 | 2 | 4 | 31 |
| | | Pontine Tegmentum | 40 | 39 | 2 | 19 | 21 | 21 | 142 |
| | | Basal Pons | 12 | 12 | 4 | 7 | 8 | 5 | 48 |
| | | Myelencephelon | 83 | 76 | 16 | 44 | 35 | 23 | 277 |
| | | White Matter | 1 | 4 | 7 | 2 | 2 | 4 | 20 |
| | | **Total** | 893 | 946 | 363 | 529 | 470 | 501 | **3702** |

Figure S1: **Anatomical organization of samples in the AHBA.**

#### S2 Topic size distribution

In the analysis, we mainly focused on the second level of the hierarchical organization of partitions given as output by our algorithm (hSBM), both for the probe layer and the sample layer. For the former, we obtained 32 topics (gene sets) at the second level of the hierarchy. After re-annotating each probe to its corresponding gene, we found that some topics have a very low number of genes, which we decided to exclude from further analysis. Below (Fig. S2), we show the distribution of the topic sizes, highlighting in red those with 10 or fewer genes, which we chose to cut.

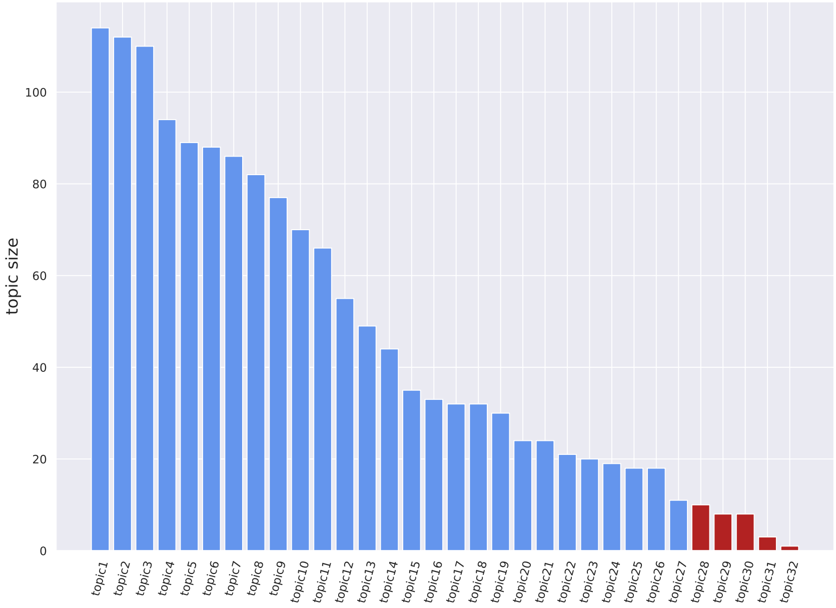


Figure S2: **Topic size distribution**. Each column is a topic from the second level of partition and the height of the column is proportional to the number of genes contained in the topic. The tail of distribution (in red) contains topics with 10 or less genes which were eliminated from the subsequent analyses.

### S3 Fuzzy membership of genes within topics

We run our algorithm directly on the probes, thus, as mentioned in the main text, we had a fuzzy memebership of genes within topics. A high percentage of genes (73%) are associated with at least two probes, and it may happen that probes of the same genes are allocated to different topics. Consequently, moving from probes to genes results in an overlap between gene topics. We report in Figure S3 the matrix of overlaps between different topics.

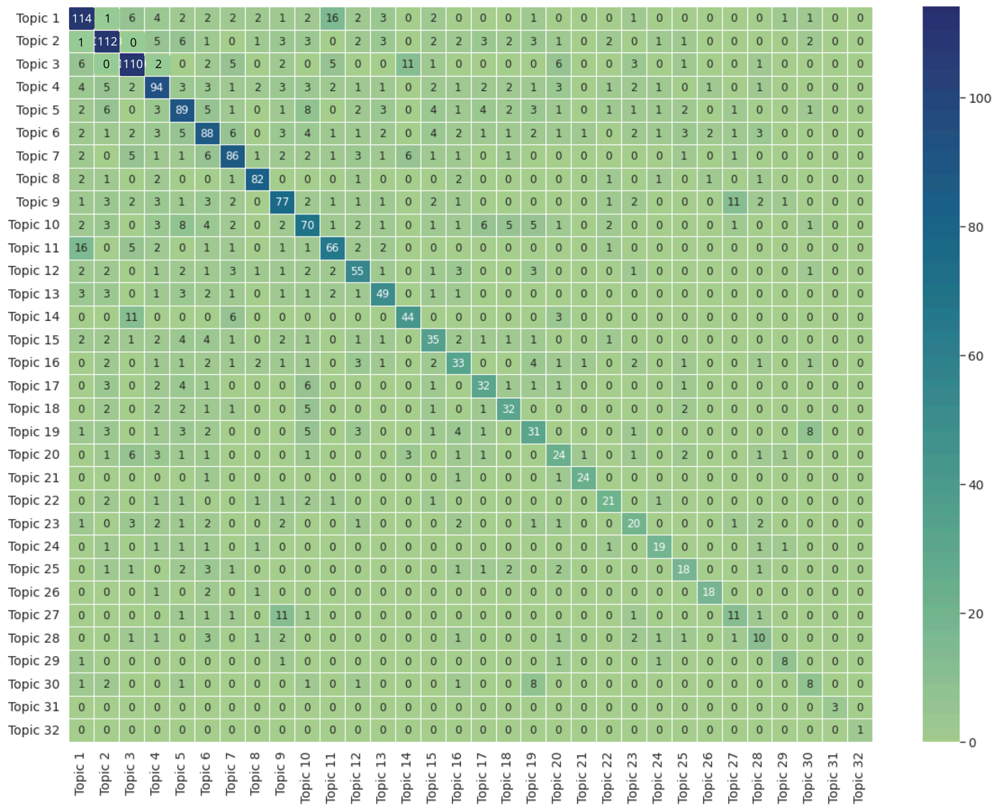


Figure S3: **Number of genes shared among gene topics.** Each matrix entry represents the number of genes common between the two topics which label the columns and the rows. The possible overlap occurs because we employ hSBM algorithm on a probes x samples matrix, resulting in gene clusterization into topics, which are sets of probes.

#### S4 NMI* scores achieved by hSBM with respect to different gene selection choices

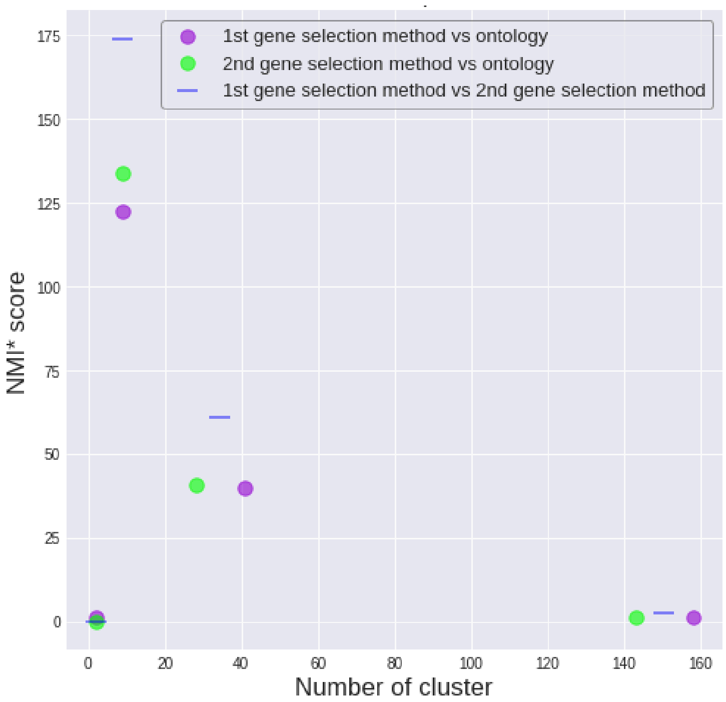


Figure S4: **Performance of hSBM with different gene selections.** We ran hSBM 10 times and averaged the corresponding NMI* scores (recall that the NMI* score quantifies the agreement between the clusters obtained with hSBM and the annotation of samples in regions and subregions). The figure reports the comparison made running hSBM with highly variable genes within the merged dataset of the six brains (1st gene selection method in the figure) or independently selecting the most highly variable genes for each brain, and keeping the intersection among them (2nd gene selection method in the figure). The scores obtained with the two selections, represented by dots of different colors, are essentially equivalent, with a NMI* score slighty higher for the second method with respect to the first one. The blue lines represent the NMI* values obtained by comparing the two partitions between them. Its very high value for the two intermediate layers shows that, despite the different gene selections the samples are clustered by hSBM in a very similar way. The first and last layers of clustering are not informative and in fact the NMI essentially coincides with the random one. The results reported in the main text (the blue dots of fig.4d) refer to the first gene selection method.

#### S5 Comparison between hSBM topics and Hawrylycz et al. modules

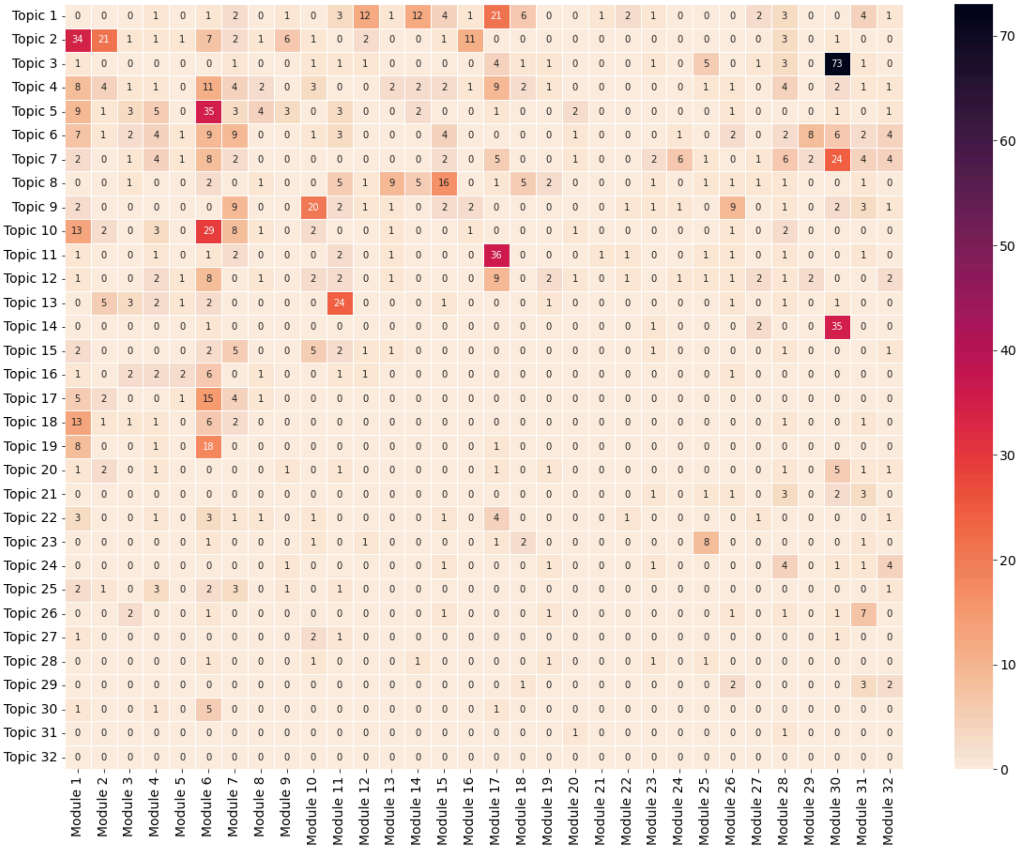


Figure S5: **Overlap between the gene content of the hSBM topics and the modules of Hawrylycz et al.** Each entry of the matrix represents the number of genes in common beteween the select topic (on the rows) and module (on the columns).

### Result Data

The following files, obtained as results of the main work, are made available in the dedicated drive folder

- *topsbm_level_2_topics_sorted.txt*: file containing genes divided into 32 topics, obtained as output from the second resolution level, by running hierarchical stochastic Block Model with the probes-samples matrix obtained by filtering highly variable probes across the merged dataset of six brains. Genes are sorted based on the probability of the gene belonging to the respective topic. For each gene, this probability was obtained by considering the highest probability among those of the probes associated with the gene.

- *topsbm_level_3_topics.csv* - *topsbm_level_4_topics.csv*: files containing probes divided into 164 (lelvel 3) and 331 (level 4) topics, obtained as output from the third and fourth resolution levels, by running hierarchical stochastic Block Model with the probes-samples matrix obtained by filtering highly variable probes across the merged dataset of six brains. Probes are not sorted.

- *topsbm_level_2_clusters.csv* - *topsbm_level_3_clusters.csv* - *topsbm_level_4_clusters.csv*: files containing tissue samples divided into 9 (lelvel 2), 41 (level 3) and 158 (level 4) clusters, obtained as output from the second, third and fourth resolution levels, by running hierarchical stochastic Block Model with the probes-samples matrix obtained by filtering highly variable probes across the merged dataset of six brains. Samples are not sorted.
